# A human iPSC-astroglia neurodevelopmental model reveals divergent transcriptomic patterns in schizophrenia

**DOI:** 10.1101/2020.11.07.372839

**Authors:** Attila Szabo, Ibrahim A. Akkouh, Matthieu Vandenberghe, Jordi Requena Osete, Timothy Hughes, Vivi Heine, Olav B. Smeland, Joel C. Glover, Ole A. Andreassen, Srdjan Djurovic

**Affiliations:** NORMENT Center of Excellence (CoE), Institute of Clinical Medicine, University of Oslo, and Division of Mental Health and Addiction, Oslo University Hospital, Oslo, Norway; Department of Medical Genetics, Oslo University Hospital, Oslo, Norway; Complex Trait Genetics, Center for Neurogenomics and Cognitive Research, Amsterdam Neuroscience, Vrije Universiteit, Amsterdam, The Netherlands; Laboratory for Neural Development and Optical Recording (NDEVOR), Section for Physiology, Department of Molecular Medicine, Institute of Basic Medical Sciences, University of Oslo, Oslo, Norway; Norwegian Center for Stem Cell Research, Department of Immunology and Transfusion Medicine, Oslo University Hospital, Oslo, Norway; NORMENT CoE, Department of Clinical Science, University of Bergen, Bergen, Norway

**Keywords:** schizophrenia, astrocyte, iPSC, bioinformatics, RNA-sequencing

## Abstract

While neurodevelopmental abnormalities have been associated with schizophrenia (SCZ), the role of astroglia in disease pathophysiology remains poorly understood. In this study we used a human induced pluripotent stem cell (iPSC)-derived astrocyte model to investigate the temporal patterns of astroglia differentiation during developmental stages critical for SCZ using RNA-sequencing. The model generated astrocyte-specific patterns of gene expression during differentiation, and demonstrated that these patterns correspond well to astroglia-specific expression signatures of *in vivo* cortical fetal development. Applying this model, we were able to identify SCZ-specific expression dynamics in human astrocytes, and found that SCZ-associated differentially expressed genes were significantly enriched in the medial prefrontal cortex, striatum, and temporal lobe, targeting *VWA5A* and *ADAMTS19*. In addition, SCZ astrocytes displayed alterations in calcium signaling, and significantly decreased glutamate uptake and metalloproteinase activity relative to controls. These results provide strong support for the validity of our astrocyte model, and implicate novel transcriptional dynamics in astrocyte differentiation in SCZ together with functional changes that are potentially important biological components of SCZ pathology.

## INTRODUCTION

Schizophrenia (SCZ) is a severe psychiatric disorder with a considerable disease burden and a lifetime prevalence of 0.3-0.7%.^1,2^ It is characterized by substantial phenotypic heterogeneity, including various cognitive dysfunctions and a profound adverse psychosocial impact. It is a highly heritable disease and well-powered genome-wide associations studies (GWAS) have identified numerous risk loci, many of which have been robustly associated with SCZ across several studies. However, the details of the disorder’s pathophysiology and etiology are still unknown,^3,4^ and no effective therapeutic modalities have been identified that could comprehensively treat the disease.^5^

To date, the neurodevelopmental hypothesis has been the dominant paradigm for the etiology of SCZ.^6^ This hypothesis posits that a significant proportion of disease vulnerability is explained by subtle changes in brain cytoarchitecture associated with structural abnormalities during the early phases of brain development.^7,8^ A combination of genetic and environmental factors is likely to play a role in disrupting the normal morphogenesis of the fetal brain. In addition, an interplay of various exogenous and endogenous components may affect CNS development and maturation, giving rise to the biological basis from which psychosis and other clinical symptoms emerge, most commonly during adolescence and early adulthood.^9,10^

Since their discovery over a century ago, astrocytes have emerged as the predominant glial cell type in the human brain involved in vital CNS functions, such as the formation of the blood brain barrier, regulation of neurotransmission and synaptogenesis, and immune and metabolic support of neurons.^11,12^ Recent human post-mortem studies have shown that alterations in astrocyte functions are associated with SCZ and other severe mental disorders, such as bipolar disorder.^13,14^ However, the role of astroglia in the pathophysiology of severe mental disorders remains poorly understood. It is therefore important to elucidate the cellular properties that influence astrocyte differentiation and maturation during brain development, as well as guide their colonization and sculpting of both the gray and white matters of the CNS in SCZ.^12,15,16^

The inaccessibility of human CNS tissue and the lack of appropriate animal models of SCZ have limited our ability to find direct evidence for astroglial involvement in SCZ brain pathology. However, recent methodological developments in human induced pluripotent stem cell (iPSC) biology has opened up novel avenues for the investigation of the cellular and molecular aspects of SCZ.^17-21^ Here, we present an *in vitro* human iPSC-astrocyte model that enables the characterization of differentiating iPSC-derived astrocytes at high temporal resolution using several data points from baseline (iPSC) to the mature astrocyte stage (day 40). We perform comparative analyses (SCZ versus healthy controls) using a combination of high coverage RNA-sequencing (RNA-seq) and astrocyte functional assays to identify possible cellular and molecular features of SCZ in developing and fully differentiated iPSC-astroglia.

## RESULTS

### Molecular characterization of differentiating iPSC-derived astroglia

To establish an *in vitro* human astrocyte model for the investigation of cellular and molecular features of SCZ, we selected one iPSC clone from each donor (5 CTRLs and 5 SCZs) at passage number 24-25. We derived astrocyte-like cells using the modified version of two previously published glial differentiation protocols,^22,23^ as we reported recently.^21^ The applied glial induction conditions successfully generated cell populations with rosette-like structures derived from embryoid bodies. Areas outside the neural rosettes exhibited varying morphologies. To evaluate the different neural progenitor (NP) populations for the expression of known markers and monitor the differentiation of astrocytes from the earliest stages (day 0 iPSCs), a custom-designed TLDA gene array card was used to assess the expression of the NP markers *PAX6* and *Nestin*, the key astrocyte markers *GFAP, S100B, AQP4, SLC1A2* and *SLC1A3*, and additional astroglia-specific markers, such as *FABP7, ALDH1L1*, and *ALDOC*.

All cultures in the neural rosette/NP stage displayed high mRNA expression of the critical markers PAX6 and NES, and these markers were continuously expressed, although progressively down-regulated over time, in the early progenitor stages and in differentiated cells (Fig. 1A-B), which is in line with previous reports.^24-26^ All fully differentiated lines manifested the typical expression kinetics of developing astroglial marker genes (*GFAP, S100B, AQP4, SLC1A2, SLC1A3, FABP7, ALDH1L1*; see Fig. 1B).^12^ Astrocyte markers that typically appear later in development (including GFAP, S100B, SLC1A2, SLC1A3, FABP7) showed robust mRNA expression only in the late, corresponding stages of differentiation (day 30 and 40; Fig. 1B), confirming an appropriate differentiation trajectory of cell cultures.^12,27,28^ Day 40 iPSC-astrocyte cultures exhibited a moderate expression of the *ALDH1L1* gene, while the mRNA expression of ALDOC was not modulated significantly during differentiation, indicating a skewing process towards cortical (protoplasmic) cellular identity in our iPSC-astrocytes cultures (Fig. 1B).^27,28^ No statistically significant differences were detected between controls and patients in the mRNA expression patterns of astrocyte-specific markers at any timepoint during the differentiation process (Fig. 1B). The astrocyte phenotype of the fully differentiated iPSC-derived cells was further validated by immunocytochemistry (ICC) demonstrating the expression of the key astroglial markers GFAP, S100B, and AQP4 in all cultures on day 40, with no observable morphological difference between SCZ astrocytes and CTRLs (Fig. 1C and Supplementary Fig. 1). In addition, our ICC results showed a good enrichment of astroglia in day 40 cultures with high purity of GFAP (>90%), S100B (>99%), and AQP4 (>96%) expressing cells (Suppl. Table 1).

**Figure 1.**
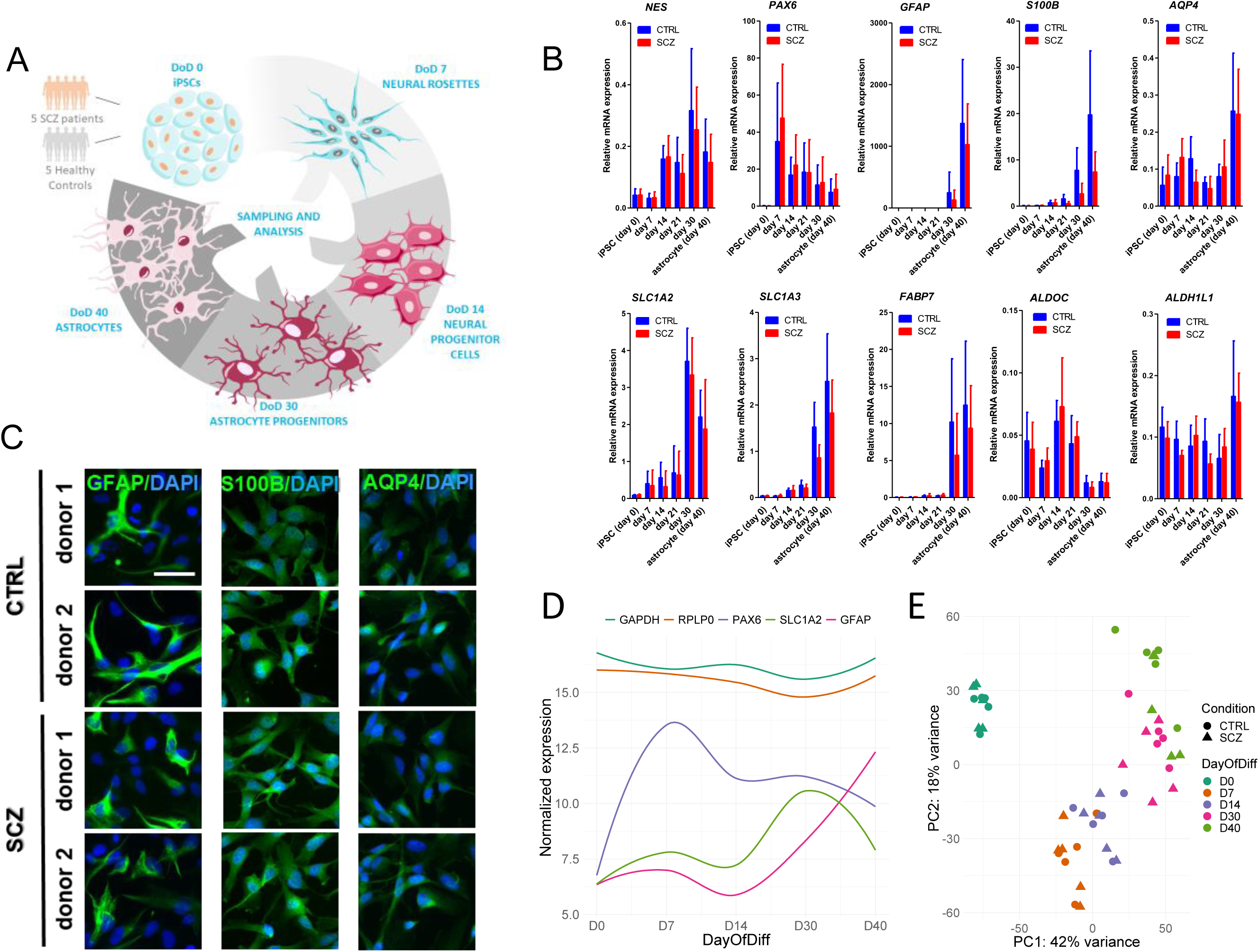
Characterization of iPSC-derived astrocytes. **(A)** Diagram of the main stages of the study. DoD: day of differentiation. This figure was created using Servier Medical Art templates, which are licensed under a Creative Commons Attribution 3.0 Unported License; https://smart.servier.com **(B)** Gene expression analysis by qPCR of NPC (PAX6, NES) and astrocyte-specific markers (GFAP, S100B, SLC1A2, SLC1A3, FABP7, AQP4, ALDH1L1, ALDOC) in SCZ (red bars) and CTRL cultures (blue bars). Data of triplicate measurements of five independent donors per group are presented as Mean ± SEM. **(C)** Cells from day 40 cultures stained positive for the key astrocyte markers GFAP, S100B, and AQP4. Images of 2 representative CTRLs and 2 donors with SCZ out of 5 are presented. Scale bar: 200 μm. **(D)** Normalized gene expression patterns of two housekeeping genes (*GAPDH* and *RPLP0*) and specific markers of astrocyte differentiation. Confidence interval: 0.5. **(E)** Principal component analysis (PCA) showing clustering of samples across the first two principal components (PC) based on the top 1000 genes with most variance between samples.

To investigate whether high-coverage RNA-seq data confirmed an astrocyte-specific pattern of gene expression across differentiation, we examined the expression levels of two housekeeping genes (*GAPDH* and *RPLP0)* and multiple markers of specific stages of astrocyte differentiation (Fig. 1D). As expected, the housekeeping genes showed a consistently high level of expression throughout. The neural rosette marker *PAX6* peaked at day 7 and continued to be expressed at slightly lower levels, while the astrocyte progenitor marker *SLC1A2* was highly expressed only at day 30. The expression of *GFAP* started to increase with the formation of progenitor cells (day 30) and reached a peak when mature astrocytes began to appear. None of the examined neural progenitor cell (NPC; day 14) markers displayed a distinctively high expression level at this stage. Next, we applied principal component (PCA) analysis to the RNA-seq data in order to explore to what extent gene expression variation is explained by the differentiation process (Fig. 1E). When clustering across the first two components, the iPSC (day 0) population separated totally from the remaining four populations. Although the separation of the other timepoints was not as complete, they nevertheless formed clear and distinct clusters, demonstrating that the variation in gene expression is mainly explained by progression through astrocyte differentiation. By contrast, no clear separation on diagnostic status (CTRL vs. SCZ) was observed.

### *In vitro* gene expression profiling of differentiating astrocytes shows similarities with existing data sets and corresponds to *in vivo* cortical fetal development

To assess the transcriptional similarity of our iPSC-derived NPCs and astrocytes to other iPSC studies, we compared our RNA-seq data from CTRL samples with the publicly available iPSC-NPC and iPSC-astrocyte, as well as primary human astrocyte expression data sets generated by Tcw et al.,^23^ the study upon which our iPSC model was partly based. Using PCA, we found that most of the variation in gene expression was due to laboratory batch effects (Fig. 2A), which was expected given that iPSC-derived differentiation protocols tend to show poor cross-site reproducibility.^29^ However, when these batch effects were adjusted for, the PCA revealed that our differentiated astrocytes were similar to the external iPSC-astrocytes and primary astrocytes, while our iPSC-derived NPCs formed a separate cluster with the external NPC samples (Fig. 2A). Hierarchical clustering showed a similar clustering pattern, with all iPSC-based astrocytes, with the exception of one sample, being more similar to each other than to the iPSC-derived NPCs, both ours and the external (Fig. 2B). Finally, correlation analyses demonstrated that the global gene expression patterns of our differentiated astrocytes were better correlated with both the external iPSC astrocytes (r=0.83) and primary astrocytes (r=0.82) than with the external iPSC NPCs (r=0.76; Fig. 2C). Taken together, these results suggest that the applied iPSC model generated astrocytes with molecular characteristics highly similar to those of other iPSC-derived astrocytes as well as primary human astrocytes.

**Figure 2.**
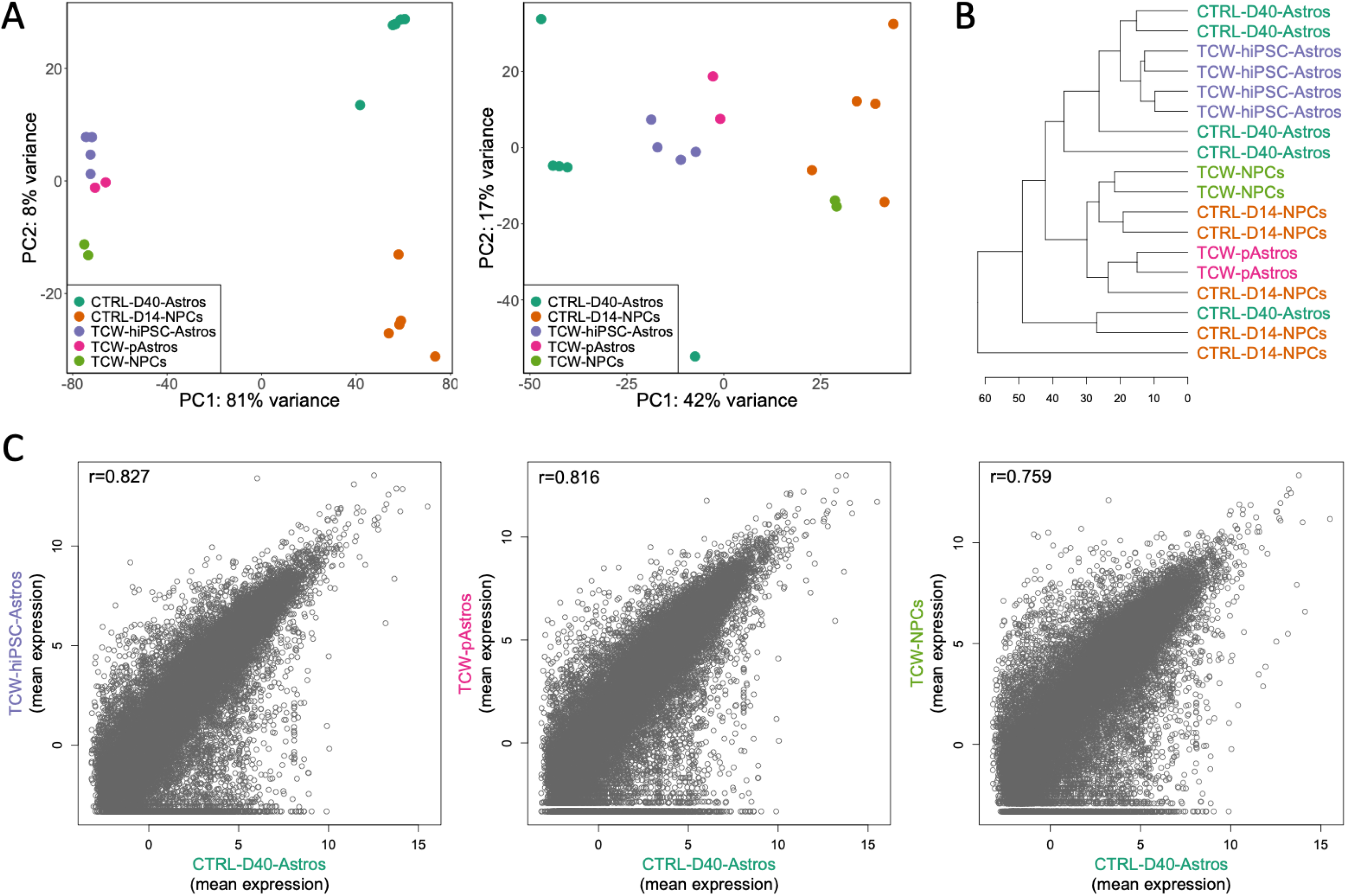
Assessment of the transcriptional similarity between iPSC-derived astrocytes and existing data sets. **(A)** PCA plots of day 14 NPCs (CTRL-D40-NPCs) and day 40 astrocytes (CTRL-D40-Astros) together with three cell types from Tcw *et al*. (see ref. 23). The left plot shows, as expected, a strong separation due to laboratory batch effects. After the laboratory batch effects were removed (right plot), the samples were well separated according to cell type across both data sets, with a relatively large variation seen within the day 40 astrocytes. TCW-NPCs: iPSC-derived NPCs, TCW-hiPSC-Astros: iPSC-derived astrocytes, TCW-pAstros: Primary astrocytes from midbrain and cerebral cortex. **(B)** Hierarchical clustering indicating good similarity between our day 14 NPCs and the TCW-NPCs, and between our day 40 astrocytes and the TCW-hiPSC-Astros, with the exception of one of our astrocyte samples which clusters together with the NPCs. Unexpectedly, the TCW-pAstros cluster together with the NPC lines. **(C)** Correlation plots of mean (normalized and log-transformed) expression data. The day 40 astrocytes show a stronger correlation with TCW-hiPSC-Astros and TCW-pAstros than with TCW-NPCs, indicating a close similarity between the astrocytes from the two studies.

Having established the astrocytic identity of the differentiated cells, we next applied transition mapping to assess whether the *in vitro* changes in gene expression during differentiation correspond to *in vivo* changes in gene expression during cortical development. While the first *in vitro* transition from iPSCs (day 0) to neural rosettes (day 7) did not match any *in vivo* transition (Suppl. Fig. 2), the formation of NPCs (day 14) and astrocyte progenitors (day 30) showed a weak correlation with the *in vivo* transition to developmental periods 24-38 PCW and 8-10 PCW, respectively (Suppl. Figs. 3-4). The most significant overlap, however, was found between the *in vitro* transition from astrocyte progenitors (day 30) to mature astrocytes (day 40) and the *in vivo* transition from 24 PCW to 38 PCW (Fig. 3), coinciding with the primary period of gliogenesis in the developing fetal cortex.^30^ These results indicate that *in vitro* differentiation of glial cells can partly capture the transcriptional dynamics of *in vivo* brain development.

**Figure 3.**
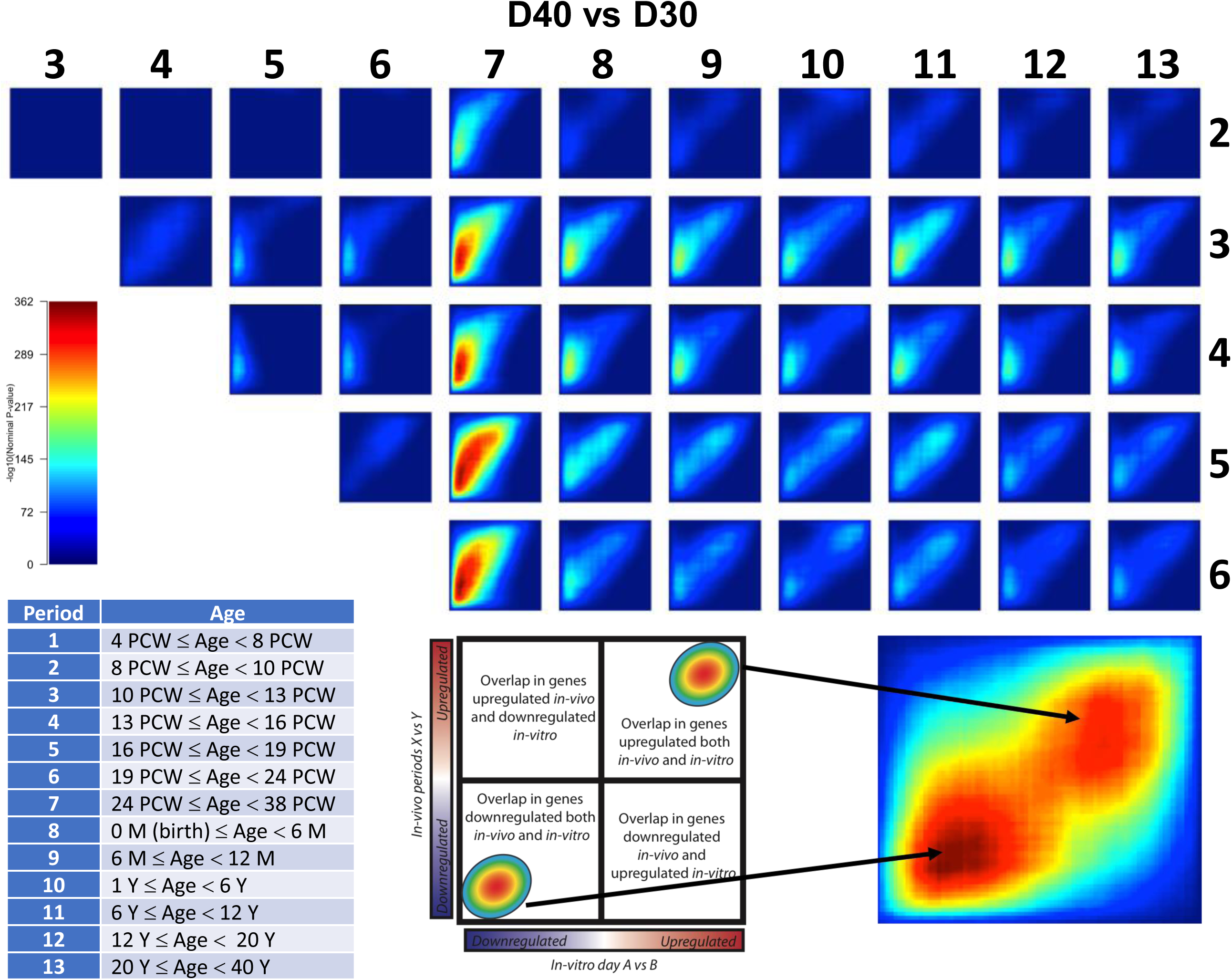
*In vitro* gene expression profiles correspond to *in vivo* cortical fetal development. Transition mapping output showing the degree of overlap between the *in vitro* transition from day 30 to day 40 (astrocytes) and *in vivo* transitions between successive periods of cortical development. The maps are colored by −log10(p-value). Strongest overlap was observed with *in vivo* transitions to period 7 (24 PCW to 38 PCW), involving mostly down-regulated genes in both *in vitro* and *in vivo* transitions. PCW: Post conception week, M: Month, Y: Year.

### Genes associated with astroglial differentiation form longitudinal clusters with relevant functions

To identify the genes that are differentially expressed (DE) during astroglial differentiation, we conducted DE analyses between successive stages along the differentiation timeline (day 7 vs. day 0, day 14 vs. day 7, etc.). In total, 9936 genes (7605 unique) were associated with astrocyte differentiation, the majority of which were related to the transition from the iPSC state (Suppl. Figs. 5A and 5C; Suppl. Table 2). Most of the identified DE genes were protein coding (66%), but a substantial proportion were long non-coding RNA genes (Suppl. Fig. 5B). Since soft clustering methods are known to be robust to the noise inherent in gene expression data,^31^ we applied fuzzy C-means clustering to the 7605 unique differentiation-DE genes and identified 7 distinct clusters consisting of genes with shared expression trajectories (Fig. 4A; Suppl. Fig. 6; Suppl. Table 3). Functional annotations of these clusters revealed that biological processes related to synaptic functions (clusters 4 and 5) were generally up-regulated as mature astrocytes started to form, while processes related to organization of extracellular structure (clusters 1, 2 and 7) were generally down-regulated as the differentiation progressed towards astrocyte generation (Fig. 4B; Suppl. Table 4).

**Figure 4.**
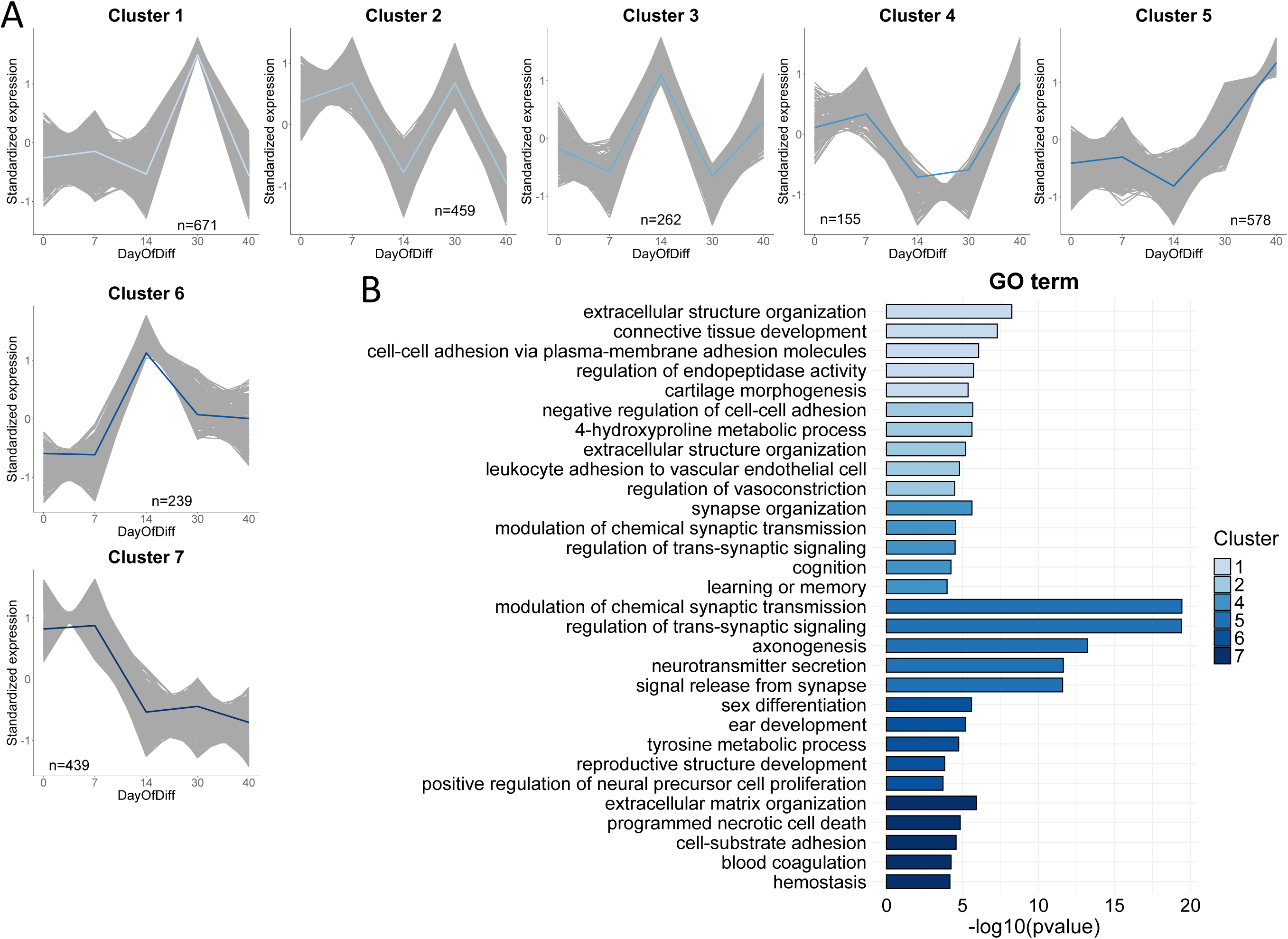
Differentiation-associated gene clusters highlight biological functions important for astroglial differentiation. **(A)** DE genes involved in astrocyte differentiation form 7 distinct clusters with shared expression trajectories. Only DE genes with cluster membership >0.5 are visualized. The number of genes in each cluster is indicated. The blue-colored lines represent cluster centroids (mean gene expression). **(B)** Top significant functional annotations (biological processes) and corresponding −log10(p-value) are shown for each gene cluster. Cluster colors correspond to the centroid colors in (A). Cluster annotations are based on DE genes with membership >0.5. Cluster 3 was not significantly enriched for any GO term. GO: Gene Ontology.

**Figure 5.**
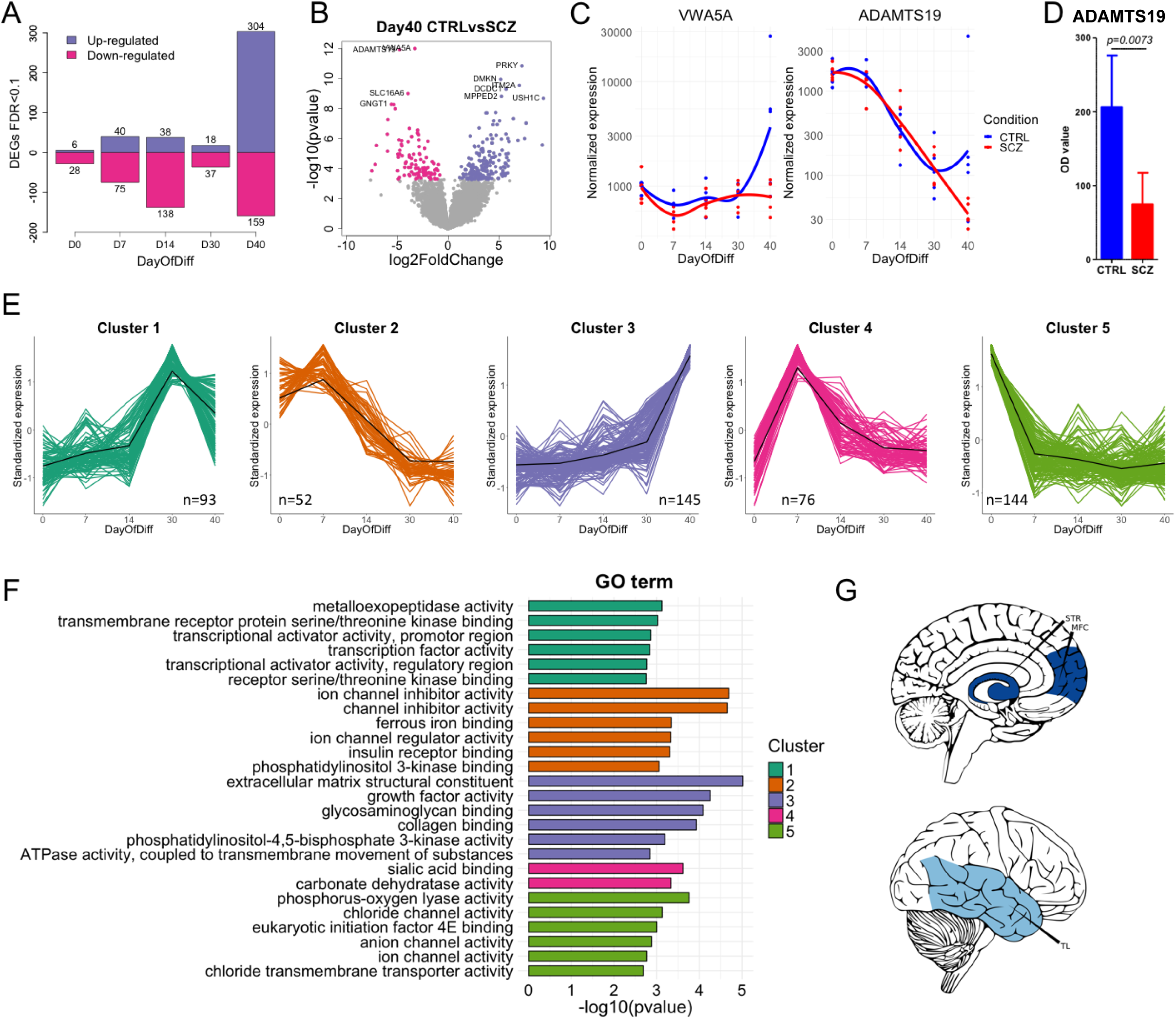
SCZ-associated genes and gene clusters are involved in ion channel and matrix metalloproteinase activity, and are enriched in specific brain regions. **(A)** Barplot of significant SCZ-related genes up and down-regulated at each stage of differentiation. **(B)** Volcano plot showing the logarithmic fold change and p-value of each gene with differential expression in day 40 SCZ astrocytes compared to CTRLs. **(C)** Normalized expression trajectory of the top two significant day 40 SCZ-associated genes. **(D)** Assessment of protein expression of ADAMTS19 in day 40 astrocyte cultures using ELISA. Data is based on triplicate measurements in cell cultures from all 10 donors, 5 CTRL (blue) and 5 SCZ (red). Relative ADAMTS19 protein expression is presented as mean OD value ± SEM. **(E)** Soft clustering of SCZ-associated DE genes resulted in 5 distinct clusters with shared expression trajectories. The number of genes in each cluster is shown. Only genes with cluster membership >0.5 were used for cluster generation. The black lines (centroids) represent mean standardized expression across SCZ samples. **(F)** Top significant cluster annotations (molecular functions) and corresponding −log10(p-value) for each gene cluster. GO: Gene Ontology. **(G)** Enrichment of SCZ-associated genes in specific brain regions in the developing human brain. Dark blue indicates brain regions with adjusted enrichment p-value <0.05. Light blue indicates brain regions with adjusted enrichment p-value=0.102. STR: Striatum, MFC: Medial prefrontal cortex, TL: Temporal lobe.

**Figure 6.**
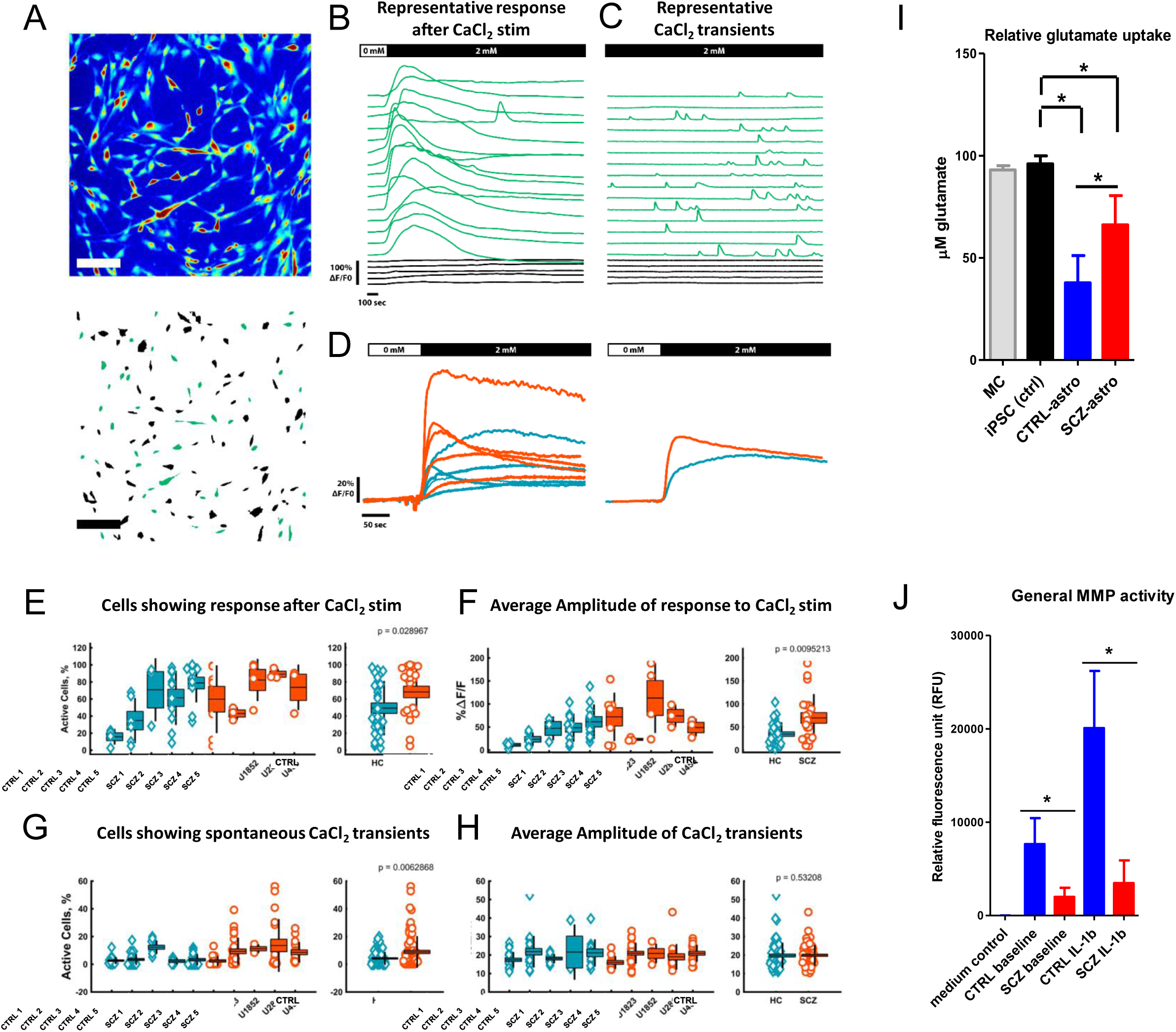
Functional characterization of hiPSC-astroglia. **(A)** Top: Representative time projection image of a 2400 frame scan obtained by live cell calcium imaging. Bottom: Representation of ROIs after segmentation, green ROIs are active cells. Scale bar equals 200μm. **(B)** Representative time course of astrocytes response to addition of 2mM CaCl_2_ in the calcium-free buffer after 60sec. Black lines are non-active cells. Green lines are active cells. **(C)** Representative time course of astrocytes’ spontaneous CaCl_2_ transients over a 20min time period. **(D)** Averaged time courses of the Calcium response of each clones (left) and, CTRL and SCZ clones compared (right). **(E)** Percentage of astrocytes with a response to the addition of 2mM CaCl_2_ in the calcium-free buffer for each clones (left), and CTRL and SCZ clones compared (right). **(F)** Average amplitude of the response to the CaCl_2_ stimulation of the active cells for each clones (left), and CTRL and SCZ clones compared (right). **(G)** Percentage of astrocytes with spontaneous CaCl_2_ transients for each clone (left), and CTRL and SCZ clones compared (right). **(H)** Average amplitude of CaCl_2_ transients for each clone (left), and CTRL and SCZ clones compared (right). **(I)** Glutamate uptake of resting iPSC-astrocytes was assessed using a glutamate assay kit. Native AM cell medium (MC, ‘medium control’), and donor/clone-specific iPSC lines were used as controls (iPSC ctrl). Data of triplicate measurements of five independent donors per group are presented as mean ± SEM (in the case of MC, triplicate measurements were performed by sampling the same bottle of medium in which all cells were cultured). Asterisks represent significant differences relative to iPSC controls, and CTRL astrocytes versus SCZ astrocytes at p < 0.05. **(J)** General matrix metalloproteinase activity was measured at both baseline and following cell activation by 10 ng/ml IL-1β for 24h. Samples from empty wells containing only AM medium were used as medium (negative) control. Fluorescence intensity was measured at E_x_/E_m_ = 540/590 nm, and MMP activity is shown in relative fluorescence units (RFU). Data of triplicate measurements of five independent donors per group are presented as mean ± SEM. Asterisks represent significant differences at p-values < 0.05.

### Genes associated with SCZ during astroglial differentiation are involved in ion channel and matrix metalloproteinase activity and enriched in specific brain regions

Next, we conducted DE analyses between CTRL and SCZ donors at each differentiation timepoint to identify genes with differential expression profiles in SCZ relative to CTRL samples. A total of 843 SCZ-associated DE genes (786 unique) were identified (Fig. 5A; Suppl. Fig. 7; Suppl. Table 5), 463 of which (55%) were differentially expressed between mature day 40 CTRL and SCZ astrocytes (Fig. 5A-B). Of these, the two most significant DE genes were *VWA5A* (FDR=1.6e-8) and *ADAMTS19* (FDR=1.6e-8), both of which were significantly down-regulated in SCZ astrocytes relative to CTRLs at day 40 (Fig. 5B-C). We measured these genes at the protein level with ELISA, which did not detect any VWA5A protein (data not shown), but did confirm significantly reduced levels of ADAMTS19 (p=0.0073) in SCZ astrocytes relative to CTRLs (Fig. 5D). Fuzzy C-means clustering of the 786 unique SCZ-related genes revealed 5 distinct clusters (Fig. 5E; Suppl. Fig. 8; Suppl. Table 6). Annotation of these clusters showed that molecular functions related to ion channel activity (clusters 2 and 5) displayed a downward trend during differentiation in SCZ patients, while functions related to transcriptional regulation (cluster 1) and extracellular matrix structure (cluster 3) displayed an upward trajectory (Fig. 5E-F; Suppl. Fig. 9; Suppl. Table 7). Cluster 4, which consisted of SCZ-associated genes involved in sialic acid binding and carbonate dehydrase activity, was first up-regulated as neural rosettes formed, and then down-regulated through the rest of the process. To examine whether the identified SCZ-related genes were over-represented in any specific brain regions, we applied a gene set expression enrichment test. The SCZ-associated DE genes were significantly enriched in the medial prefrontal cortex (MFC, FDR<0.05) and striatum (STR, FDR<0.05), and somewhat less enriched in the temporal lobe (TL, FDR=0.102; Fig. 5G), suggesting that astrocyte differentiation in these brain areas of SCZ patients may be affected.

### Functional characterization of iPSC-astroglia reveals altered cell-intrinsic traits in SCZ

The *in vivo* role of astrocyte calcium signaling is fundamental to the astroglia network and to normal brain physiology. Since our previous results showed differential expression of genes related to ion channel activity in SCZ versus CTRLs, we next examined possible discrepancies between SCZ and CTRL iPSC-astrocyte Ca^2+^ transients. Increases of astrocyte intracellular Ca^2+^ concentrations can be evoked by extracellular stimuli, such as glutamate or various other compounds (ATP, calcium salts, etc.). The subsequent propagation of Ca^2+^ signals along astroglial processes and between glial cells have been suggested to modulate neuronal functions either directly or via the regulation of gliotransmitter release.^32-36^ Activated astrocytes generate intracellular Ca^2+^ transients that are spatiotemporally ‘fine-tuned’ with reduced frequency in the cell somata relative to the processes.^37-39^ We used live fluorescence imaging in combination with high-precision Ca^2+-^indicators in single cells to characterize the functionality of iPSC-astrocytes (Fig. 6A-H).

After demonstrating that all the iPSC-astroglia lines show both spontaneous calcium transients and CaCl_2_-evoked calcium signals (Fig. 6A-D), we found that SCZ cultures were characterized by significantly higher number of active astrocytes and a more rapid and stronger calcium-evoked calcium response (Fig. 6E-H). We next sought to confirm glutamate uptake by the cells which is another important functional feature of astrocytes.^40^ Since ADAMTS19 was differentially expressed in SCZ astrocytes versus CTRLs (Fig. 5C-D) and astrocyte metalloproteinases have previously been linked to altered synaptic glutamate transport,^41,42^ we hypothesized that SCZ and CTRL astrocytes will present dissimilarities with regards to their capacity to internalize glutamate from the environment in an *in vitro* model. We found that SCZ astrocytes exhibited an approximately two-fold lower glutamate-uptake relative to healthy controls (Fig. 6I).

The brain extracellular matrix (EM) is a critical element in the regulation of neural plasticity and network development. Since glia-derived metalloproteinases, such as ADAMTS family members, have been shown to modulate brain EM structuring through the cleaving and reorganization of synaptic architecture in both the early postnatal period and in the adult brain,^43,44^ we tested the general metalloproteinase activity in iPSC-astrocytes. We observed a greatly reduced general metalloproteinase activity in SCZ astrocytes relative to controls both at baseline (∼3-fold), and following IL-1β stimulation (∼4.5-fold, Fig. 6J), a known inducer of ADAMTS family protein expression and function.^45,46^ This suggest a decreased capability of SCZ astrocytes in regulating neuroplasticity in the human brain *in vivo*, perhaps in the areas identified by our RNA-seq measurements (Fig. 5G).

## DISCUSSION

In this study we used an iPSC-astrocyte developmental model consisting of five time points between the baseline (iPSC) and fully differentiated astroglia stages in combination with high coverage RNA-seq and functional assays. Applying this model, we show divergent transcriptomic patterns of genes involved in staging of neurodevelopment processes in SCZ iPSC-astrocytes. Many of the identified DE genes involve functions characteristic of astrocytes, such as the regulation of metalloexopeptidase and ion channel activity, EM structure reorganization, modulation of chemical synaptic transmission, axonogenesis, and signal release from synapses. The temporal representation of this divergence became more prominent in the later stages of *in vitro* development as cells gradually differentiated from astrocyte progenitors (*in vitro* day 30) to the mature astrocyte state (*in vitro* day 40). Our findings on the underlying patterns in gene expression and cellular behavior that characterize several SCZ-specific alterations in human astrocyte functions provide novel insights into the neurodevelopmental pathology involved in SCZ.

All the derived cell lines used in this study exhibited the typical expression kinetics of developing astroglial marker genes. Importantly, we did not detect statistically significant differences between controls and patients in the mRNA expression patterns of the monitored astrocyte-specific markers at any time point during the differentiation process. Neither did we observe any clear morphological differences between mature SCZ astrocytes and CTRLs using confocal imaging. This suggests no apparent structural abnormalities in SCZ iPSC-astroglia and is in accordance with human post-mortem studies which report differences in astrocyte numbers^47,48^ and density in certain brain regions,^49-51^ but not in cellular morphology.^52^

Hypotheses concerning the etiology of SCZ have been greatly affected by early studies suggesting an important role for neurodevelopmental effects in the prenatal period.^6^ The dominant developmental hypothesis postulates that a considerable part of disease vulnerability is based on pathological alterations in certain brain regions during the early phases of development.^7,8^ To investigate the possible involvement of astrocytes in this process, we performed high-resolution temporal screening of differentiating iPSC-astrocyte cultures using RNA-seq gene expression profiling. These results confirmed astrocyte-specific patterns of gene expression across differentiation and also corroborated astrocyte-specific differentiation that corresponded to *in vivo* cortical fetal development in all cultures. Transition mapping results revealed that *in vitro* differentiation signatures corresponded to *in vivo* gene expression patterns of cortical development in a differentiation stage-dependent manner. The first correlations were seen during the *in vitro* transition from NPC (day 14) to astrocyte progenitor (day 30) stages demonstrating weak associations with the early *in vivo* developmental periods. However, a strong overlap was seen between the *in vitro* transition from astrocyte progenitors to mature astrocytes (day 40) and the *in vivo* transition from 24 PCW to 38 PCW, which is in agreement with neurodevelopmental models defining this stage as the primary period of gliogenesis in the developing fetal cortex.^30^ These results suggest that *in vitro* temporal screening of differentiating glial cells can allow us to make mechanistic inferences about the transcriptional fluctuations of the *in vivo* developing human brain.

Next, we performed DE analyses to identify the genes that are differentially expressed during *in vitro* astrocyte differentiation. We identified several distinct clusters consisting of genes with shared expression trajectories, and the subsequent functional annotations of these clusters revealed typical tendencies in transcriptional dynamics. These data, in combination with the transition mapping results, confirmed a successful *in vitro* modeling of human fetal gliogenesis and astrocyte development in both SCZ and CTRL iPSC-astroglia cultures, in accordance with the literature^30^ and with publicly available human *in vivo* transcriptomic data sets.^53^

A growing body of evidence implicates astrocyte-related abnormalities in the development of human cerebral cortex and subcortical regions in neuropsychiatric disorders.^54^ Many of these glial abnormalities have been suggested to influence neural differentiation and maturation during brain development, as well as impact various aspects of neuron-glia interactions.^12,15,16^ To investigate the potential role of astrocyte-associated factors in SCZ etiology, we sought to identify genes with differential expression profiles in SCZ relative to CTRL. We performed DE analyses between CTRL and SCZ donors at each time point across the *in vitro* differentiation process. We found that the two most significant DE genes were *VWA5A* and *ADAMTS19*, both of which were strongly down-regulated in SCZ astrocytes relative to CTRLs at day 40. In follow-up experiments, we confirmed the significantly reduced expression of ADAMTS19 protein in SCZ astrocytes relative to CTRLs (∼2.5-fold difference). We could not detect VWA5A protein in any cells, but this is consistent with the fact that VWA5A protein appears to be expressed only under immune activation but not in resting astroglia.^55^

Annotation of the clusters, yielded by fuzzy C-means clustering of the 786 unique SCZ-related gene inputs, resulted in three distinct trends. First, molecular functions related to ion channel activity were down-regulated during astroglia differentiation in SCZ. Second, biological functions related to transcriptional regulation and EM structure displayed an upward expression trajectory during SCZ astrocyte-differentiation. The third trend showed an early up-regulation of SCZ-associated genes in the neural rosette stage followed by a down-regulation tendency through the rest of the differentiation process, and involved factors related to acid-base balance regulation (carbonate dehydratase activity) and innate immunity (sialic acid binding). Interestingly, out of these SCZ-specific transcriptional trends, ion channel activity and EM structure-related patterns were the most significant. To examine whether the identified SCZ-related genes were over-represented in any specific brain regions, we applied a gene set expression enrichment test. The SCZ-associated DE genes were significantly enriched in the medial prefrontal cortex (MFC) and striatum, and displayed a weaker enrichment in the temporal lobe suggesting that astrocyte differentiation in these brain areas are affected in the developing brains of SCZ patients. The complex nature of neuropsychiatric diseases entails that their molecular substrates are subject to tight spatial regulation and possess specific expression patterns across the brain parenchyma, so that they disrupt specific cognitive functions but preserve the basic functions and most of the higher-order processes of the brain. Ion channel proteins in the human brain are regulated by diverse factors ranging from a multitude of discrete isoforms and receptor subunit combinations within ion channel families and subclasses, each of which display a specific distribution that is developmentally and spatiotemporally controlled in the brain.^56-58^ Our results demonstrating that developing iPSC-astrocytes in SCZ abnormally down-regulate ion channel-activity-related genes are in agreement with a previous report showing reduced potassium uptake in SCZ iPSC-astrocytes relative to controls,^59^ and may add to the existing literature suggesting that common variation at ion channel genes can explain some of the heritability of schizophrenia.^56,60^ Similarly, EM abnormalities^61^ and dysregulated transcription factor-level alterations in iPSC-astroglia in SCZ^59^ have been reported previously, suggesting that these factors may contribute to the pathophysiology of the disorder involving disruptions in neuronal functioning, neuron-glia communication, and aberrant glutamatergic, dopaminergic, and GABAergic neurotransmission.^61^ Our iPSC-astroglia results showing a significant enrichment of SCZ-associated DE genes in the prefrontal cortex and striatum, as well as a weaker enrichment in the temporal lobe, are in good agreement with previous studies indicating the involvement of these brain regions in the symptoms and pathogenesis of SCZ.^62-66^ Specifically, alterations in astrocyte-associated markers have been shown in the prefrontal^67,68^ and subcortical regions^69^ of the brain in SCZ. Moreover, astrocytic loss/dysfunction in the MFC has been linked to cognitive impairment in rats, suggesting a pivotal role for astroglia in normal MFC functioning in mammals.^70^ In a recent study, the authors found that many of the genes previously identified as differentially expressed in SCZ displayed high correlations with the expression profiles of astrocytes or parvalbumin interneurons.^71^ In sum, our data may reveal hitherto unknown genetic patterns in astrocyte differentiation as potentially important biological components related to the regulation of human cortical development and cognition.

Astrocyte calcium signaling is essential in normal brain physiology and is regarded as a critical *in vivo* functional aspect of human brain astroglia.^36^ Our initial results showed a differential expression of genes related to ion channel activity in SCZ versus CTRLs, and thus we performed single-cell live fluorescence imaging to characterize the functionality of our iPSC-astrocytes and to look into possible alterations in baseline and experimentally evoked Ca^2+^ signals in SCZ versus CTRL cell cultures. We showed that all the iPSC-astroglia lines exhibited both spontaneous Ca^2+^ transients and CaCl_2_-evoked calcium signals. Interestingly, we found that significantly higher number of SCZ astrocytes were spontaneously active and had calcium-evoked calcium response compared to control astrocytes. Astrocyte Ca^2+^ elevations have been shown to regulate extracellular glutamate levels *in vivo*, which consequently modulate neuronal rhythms in the cortex and trigger synchronized neuronal firing associated with different behavioral states suggesting an important regulatory role for astroglia in cortical circuits.^72-74^ Moreover, evoked astrocyte Ca^2+^ signals have been shown to induce glutamate release in the hippocampus that consequently facilitates synaptic transmission by the activation of different glutamate receptor subtypes.^75^ The role of this mechanism was also demonstrated in long-term potentiation of hippocampal CA1 pyramidal cells,^76^ suggesting the involvement of astrocytic Ca^2+^-coupled glutamate signals in learning and cognition. We confirmed glutamate uptake by our iPSC-astrocytes, an important functional feature of brain astroglia in regulating cognition and behavior.^40^ Since we detected altered astrocyte Ca^2+^ signals in SCZ cells and since the metalloproteinase ADAMTS19 was differently expressed in SCZ astrocytes versus CTRLs, coupled with the fact that astroglial metalloproteinases have previously been linked to altered synaptic glutamate transport,^41,42^ we hypothesized that SCZ and CTRL astrocytes will present dissimilarities in their glutamate uptake capacity. Indeed, we found significant difference between SCZ and CTRL astroglia in their uptake capability as SCZ astrocytes exhibited an approximately two-fold decrease in glutamate-uptake relative to healthy controls. Interestingly, we observed this lower glutamate uptake despite the SCZ astrocytes having an increased Ca^2+^ signaling. As no significant differences were found in the expression of glutamate transporter genes (*SLC1A2, SLC1A3*) in SCZ versus CTRLs at any time point of differentiation, but a decrease was observed in ion channel activity-related gene expression in SCZ astroglia, this phenomenon may be explained by the effort of SCZ astrocytes to ramp up Ca^2+^ signals in order to compensate for the aberrant/dysregulated glutamate transporter functions. According to the “glutamate hypothesis of schizophrenia”, the symptoms of SCZ are rooted in NMDAR glutamate receptor hypo-function and excessive glutamate release in the synaptic cleft of cortical neurons, especially in the prefrontal cortex and hippocampus.^77^ However, it is still unknown how this NMDAR hypo-function can lead to increased glutamate and an “overdrive” of non-NMDA glutamate receptor activation. Excessive extracellular glutamate levels have been well documented in previous brain imaging studies in SCZ.^78^ Moreover, this has been shown to directly cause metabolic damage, synaptic degeneration, and cortical atrophy in the affected brain areas, and these pathological changes are highly correlated with disease symptom evolution from the prodromal phase to full-blown psychosis.^79^ Astrocytes regulate glutamatergic transmission and synaptic glutamate levels in multiple ways, ranging from glutamate biosynthesis, uptake, and release to the modulation of glutamatergic synapses via gliotransmitters, such as D-serine.^80^ Based on our results, we hypothesize that the impaired capability of SCZ astrocytes to uptake glutamate may contribute to the increased extracellular glutamate levels documented in the disease. However, more research is needed to elucidate the consequences of this phenomenon in cortical neurons and/or neuronal networks *in vitro* and *in vivo*.

The role of brain EM is critical in normal CNS development and in the regulation of neuroplasticity. Glia-derived metalloproteinases are involved in brain EM restructuring via the cleaving and shaping of synaptic architecture of both the adult and the early postnatal brain.^43,44^ We observed that SCZ astrocytes had significantly lower levels of ADAMTS19 mRNA and protein relative to CTRLs. These findings corresponded well with a strongly reduced general metalloproteinase activity in SCZ astrocytes relative to controls both at baseline and following inflammatory activation, a known inducer of ADAMTS family enzyme expression and activity.^45,46^ Since the exact biological role of ADAMTS19 is not known yet, but ADAMTS family metalloproteinases play an important role in controlling neuroplasticity and neural regeneration in the CNS,^81^ our findings may imply a decreased capability of SCZ astrocytes in regulating neuroplasticity *in vivo* in the human brain, especially in the areas revealed by our RNA-seq results (MFC, striatum, and TL).

Taken together, we report transcriptome-level differences in differentiating human iPSC-astrocytes at high temporal resolution. Our results confirm astrocyte-specific patterns of gene expression during differentiation and show that our *in vitro* astroglia differentiation method adequately model cortical fetal development. Furthermore, we also demonstrate that the *in vitro* differentiation signatures of developing iPSC-astrocytes correspond to *in vivo* gene expression patterns of cortical development in a differentiation stage-dependent manner. Our results reveal for the first time SCZ-specific expression dynamics in astroglia, demonstrate that SCZ-associated DE genes are significantly enriched in distinct brain regions, and identify *VWA5A* and *ADAMTS19* as differentially expressed genes in SCZ versus CTRLs. In corresponding functional assays we identify several abnormalities in SCZ astrocytes, such as alterations in Ca^2+^ signaling, and significantly decreased glutamate uptake and general metalloproteinase activity. In sum, these data may reveal hitherto unknown transcriptional dynamics and patterns in astrocyte differentiation, as well as SCZ-specific functional changes as potentially important biological components in disease etiology and pathology.

## MATERIALS AND METHODS

### Recruitment of patients and collection of skin biopsies

This project is part of the ongoing Norwegian TOP (Thematically Organized Psychosis) study. Information about recruitment procedures, inclusion and exclusion criteria, and clinical assessments for the TOP study as a whole have previously been described in detail.^82,83^ For reprogramming and astrocyte differentiation, skin biopsies/fibroblasts were isolated from 5 healthy controls (CTRL) and 5 SCZ patients that were selected based on clinical information (Fig. 1A). Supplementary Table 8 shows further details about the study participants from whom skin biopsies were obtained. All patients underwent a clinical examination that included diagnostic interviews based on Structured Clinical Interview in DSM-IV axis I Disorders (SCID-1) and structured assessments of clinical symptoms. Diagnostic evaluation was performed by trained clinical psychologists and psychiatrists. The main inclusion criteria were confirmed diagnosis of SCZ according to the Diagnostic and Statistical manual of Mental Disorders (DSM)-IV, age between 18-65, and ability to give informed written consent. The main exclusion criteria were clinically significant brain injury, neurological disorder, ongoing infections, autoimmune disorders or any form of cancer. All participants have given written consent and the study was approved by the Norwegian Data Protection Agency and the Regional Ethics Committee of the South-Eastern Norway Regional Health Authority (REK grant: #2012/2204). The authors assert that all procedures contributing to this work comply with the ethical standards of relevant guidelines and regulations.

### Generation of human iPSCs and differentiation of astrocytes

The generation and characterization of iPSCs and differentiation of iPSC-derived astrocytes were done as described previously.^21^ Briefly, fibroblasts isolated from control and patient donors were reprogrammed using Sendai virus, transduced with the CytoTune™-iPS 2.0 Sendai Reprogramming Kit (Thermo Fisher, Waltham, MA, USA) containing KOS (Klf4, Oct4, Sox2), Nanog, and c-Myc reprogramming factors. Each iPSC line was subjected to rigorous quality control at The Norwegian Core Facility for Human Pluripotent Stem Cells at the Norwegian Center for Stem Cell Research including phenotyping, regular monitoring of morphology, and pluripotency marker expressions. Karyotyping to assess the chromosomal integrity of all the derived iPSC lines was performed at passage 15 for verification of authenticity and normality (KaryoStat Karyotyping Service, Thermo Fisher). Karyotyping test results and iPSC phenotyping data have been published previously,^21^ and are available upon request.

Astrocyte differentiation was performed following two previously published glial differentiation protocols with moderate modifications,^22,23^ as reported elsewhere.^21^ We previously derived and characterized astrocytes from 3 CTRLs and 3 SCZ patients, and here we performed the same level of rigorous characterization and phenotyping of two additional SCZ and CTRL donor lines to increase data density (Figure 1A-C; Supplementary Figure 1). In brief, iPSC colonies at passages 24-25 were transferred to 6-well tissue culture plates in Neural Maintenance Medium (NMM) consisting of 50% DMEM/F12 and 50% Neurobasal Medium (both from Thermo Fisher) supplemented with 0.5% (v/v) N2, 1% (v/v) B27 (both from Invitrogen, Carlsbad, CA, USA), 5 μg/ml human insulin, 40 ng/ml triiodothyronine (T3), 10 μM β-mercaptoethanol, 1.5 mM L-glutamine, 100 μM NEAA, 100 U/ml penicillin and 100 μg/ml streptomycin (all from Sigma-Aldrich, St. Louis, MO, USA). 20 ng/ml EGF and 4 ng/ml bFGF (both from Peprotech, Rocky Hill, NJ, USA) were added to the cultures. After 1 day, non-adherent embryoid bodies (EBs) formed in the cultures and were fed daily with NMM containing T3, EGF and bFGF (as above) until day 3. On day 3, 10 μM all-trans retinoic acid (ATRA, Sigma) was added to the medium and EBs were washed (1x, gently in NMM medium) and plated on Geltrex-coated plates (Thermo Fisher). After this step, cultures were fed with ATRA + T3 + growth hormone supplemented NMM medium daily until day 10. On day 10, neurorosette formation was monitored by light microscopy, and *NES* (Nestin) and *PAX6* positivity was checked by qPCR (Fig. 1B). Cultures were passaged using the cell dissociation agent Accutase (Sigma) following the recommended protocol. From this point on, regular passaging was done at confluence and cultures were seeded on Geltrex-coated 6-well plates. On day 10, the medium was changed to NMM + 40ng/ml T3 + 20ng/ml EGF. On day 18, after the formation of neural progenitor cells (NPCs), the medium was changed to NMM supplemented with B27 without vitamin A (Invitrogen) + 20 ng/ml EGF + 40ng/ml T3, and cultures were fed using the same medium composition until day 40. Samples were continuously collected during the differentiation process at days 0 (iPSC), 7, 14, 21, 30, and 40. The mRNA expression of the neural progenitor markers *PAX6* and *NES*, and the astrocyte-specific markers *GFAP* (Glial fibrillary acidic protein), *S100B* (S100 calcium-binding protein B), *AQP4* (Aquaporin 4), *SLC1A2* and *SLC1A3* (Solute Carrier Family 1 Member 2 and 3), *FABP7* (Fatty acid binding protein 7), *ALDH1L1* (Aldehyde Dehydrogenase 1 Family Member L1), and *ALDOC* (Aldolase C) was monitored by qPCR using a custom-designed TLDA gene array card (Thermo Fisher). Cells were stained with anti-GFAP, anti-S100B, and anti-AQP4 antibodies (all from Abcam, Cambridge, UK) and were analyzed using fluorescence microscopy on day 40. Only cell lines displaying a successful iPSC-to-astroglia differentiation trajectory (one per donor, based on marker expression) were selected for developmental characterization by submitting samples from the main stages of *in vitro* human astrocyte development for RNA-seq (day 0, 7, 14, 30, and 40 samples; Fig 1A-C). Fully differentiated iPSC-derived astrocytes expressing the relevant astroglia-specific markers were used for functional studies on day 40.

### Immunocytochemistry (ICC) and imaging

Samples were fixed with 4% paraformaldehyde for 10 minutes, and washed 3 times with PBS for 15 minutes. Samples were blocked for 1 h with 10% normal horse serum (NHS; Abcam) in Tris-buffered saline containing 0.5% (v/v) Triton X-100, pH 7.4 (TBSTx). Fixed cells were further incubated overnight in 1% NHS in TBSTx with rabbit anti-GFAP (diluted 1:500; Dako-Agilent, Santa Clara, CA, USA), rabbit anti-S100B (1:400) or rabbit anti-AQP4 (1:500) antibodies (both from Abcam). Samples were then rinsed 3x with 1% NHS in TBSTx and blocked again for one hour at RT. Samples were incubated with Alexa Fluor 488 goat anti-rabbit IgG secondary antibody (1:500) for 1 hour at RT (Invitrogen). The samples were then rinsed 3x with 1% NHS in TBSTx and incubated for 5 min with the nuclear stain DAPI (1 μg/ml in PBS; Sigma). The samples were then rinsed 2x with PBS and mounted under coverslips in PBS:glycerol 1:1. The samples were examined and imaged with a Zeiss LSM 700 confocal microscope. Images were processed using ImageJ software version 1.52p (National Institutes of Health, USA). During culture purity assessment, the number of GFAP, S100B, or AQP4-positive astrocytes was obtained by automated cell counting using an ImageJ macro as previously described.^84^

### RNA extraction and qPCR/TLDA array

Total RNA was extracted from all samples (5 CTRL and 5 SCZ subjects at six timepoints across astrocyte differentiation) using either the RNeasy Plus Mini Kit (Qiagen, Hilden, Germany) or the MagMax mirVana Total RNA Isolation Kit (Thermo Fisher). Approximately 1 million cells were used as input. All samples had an RNA Integrity Number (RIN) above 8.5 with three exceptions: two samples at 7.8 and one sample at 6.6. cDNA was generated from 500 ng of total RNA, using the High-Capacity cDNA Reverse Transcription Kit (Life Technologies Corporation, Carlsbad, CA, USA) according to the manufacturer’s protocol. qPCR was performed with custom designed TaqMan low density array (TLDA) cards (Life Technologies; catalog numbers of the included TaqMan probes are available upon request). Briefly, 100 ng of cDNA was loaded per port according to the manufacturer’s protocol. The cards were then cycled and analyzed on a Quantstudio 12K Flex Real-Time PCR system (Life Technologies), using the manufacturers standard temperature profile. Relative target gene expression was computed with the delta-delta-Cq method in the qBasePLUS software (v3.1).

### RNA sequencing and data processing

Sequencing libraries for the 50 samples (collected on days 0, 7, 14, 30, and 40 from each of the 10 donors) were prepared with the TruSeq Stranded mRNA kit from Illumina (San Diego, CA, USA) and sequenced on a HiSeq 4000 sequencer (Illumina) at an average depth of 50 million reads per sample using a read length of 150 base pairs and an insert size of 350 base pairs. Raw sequencing reads were quality assessed with FastQC (Babraham Institute, Cambridge, UK) and further processed with Trimmomatic V0.32 using default parameters.^85^ HISAT2^86^ was then used to map the trimmed reads to the human GRCh38 reference genome. Finally, gene expression levels were quantified by summarizing the mapped reads at the gene level using featureCounts.^87^ To assess the extent to which our iPSC-derived CTRL NPCs and astrocytes resemble the cells generated by Tcw et al.,^23^ raw RNA-seq data files were downloaded from the Gene Expression Omnibus site (https://www.ncbi.nlm.nih.gov/geo/; accession number: GSE97904) and processed as described above.

### Differential expression analysis

Differential expression (DE) analyses were performed with *DESeq2*^88^ after filtering out lowly expressed genes, defined as <3 read counts in more than 50% of the samples. Two sets of DE analyses were carried out. The first set of DE analyses compared two successive stages of astroglial differentiation (day 7 vs. day 0, day 14 vs. day 7 etc.) and aimed to identify genes whose expression profiles change across the differentiation process (differentiation-associated genes). The second set of DE analyses compared CTRL subjects to SCZ patients at each stage of astrocyte differentiation in order to identify genes that were differentially expressed in SCZ during differentiation (SCZ-associated genes). Details about the statistical design and significance criteria used can be found in the Supplementary Methods.

### Transition mapping to cortical fetal development

To investigate how well the *in vitro* gene expression profiles across astrocyte differentiation correspond to *in vivo* expression patterns of the developing fetal brain, we applied transition mapping (TMAP)^30^ to the differentiation-associated genes. The *in vivo* developmental periods were defined from a publicly available spatiotemporal transcriptomic atlas.^53^ Since TMAP was originally developed for time course data from microarrays, a modified version tailored for RNA-seq was used (https://github.com/dhglab/RRHO_RNAseq).

### Fuzzy C-means clustering and cluster annotation

In order to cluster the differentiation-associated and the SCZ-associated DE genes according to their expression trajectory across astrocyte differentiation, we applied fuzzy C-means clustering to the two gene sets separately using the R package *e1071*. Fuzzy C-means clustering is a soft clustering method which is particular robust to the noise inherent in gene expression data, and it has been successfully been applied to identify clusters in time series gene expression experiments.^31^ As input, we used normalized expression values that were averaged either across all 10 samples (for differentiated-associated genes) or across SCZ samples (for SCZ-related genes) for each stage of differentiation and then standardized. Only DE genes with membership >0.5 were used to generate the cluster plots and to annotate the clusters. Cluster annotation was performed using the over-representation analysis tool *clusterProfiler*.^89^ Annotations were based on Gene Ontology (GO) terms,^90^ and a GO term was considered significantly over-represented if the FDR was <0.10.

### Enrichment in human brain regions

To test whether the SCZ-specific DE genes were enriched in any specific brain region, we used *ABAEnrichment*.^91^ As cut-off for the annotation of genes to brain regions, the default 10%-steps of expression quantiles across all regions was used. Moreover, the default “5_stages” dataset, which uses data from five developmental stages from the BrainSpan Atlas of the Developing Human Brain (http://brainspan.org/), was selected for the enrichment analyses. Enriched brain areas were visualized with *cerebroViz*.^92^

### ELISA assays

We quantified vWA5A and ADAMTS19 protein levels by lysing 5 x 10^5^ cells using Pierce IP Lysis Buffer (Thermo Fisher) following the recommended protocol and measured human vWA5A and ADAMTS19 with high-sensitivity ELISA kits (both from MyBioSource, San Diego, CA, USA; intra-assay variation: CV<10%; inter-assay variation: CV<12%).

### Astrocyte live cell calcium imaging

Cells were loaded with 1 μM Fluo-4-AM (Thermo Fisher) in culture medium and incubated at 37 °C and 5% CO_2_ for 30 minutes. Cultures were then washed using a buffer containing 1 mM MgCl_2_, 2 mM KCl, 135 mM NaCl, 1 mM MgSO_4_.7H_2_O, 10 mM HEPES, and 10 mM D-glucose. Images were recorded with an upright microscope (Axioskop FS Carl Zeiss) using an Epiplan-NEOFLUAR 10x objective lens (NA 0.25), combined with the Andor iXon Ultra 897 camera and Andor Solis software. Frame scans were acquired at 4Hz (512×512 pixels) for a period of 20 min. For each well, the first recording was performed in a CaCl_2_ free solution (as described above) representing the baseline fluorescence, then after 1 min 2mM CaCl_2_ was added to the cells. After that, 3 to 4 more recordings of 20 min were done in the CalCl_2_ containing buffer.

A custom-made MATLAB script was used to analyze live cell calcium traces and derive various parameters reflecting characteristics of astrocyte activity. In brief, regions of interest (ROIs) were selected through automatic segmentation using extended pixel representation based on time projection images of the recordings. For each ROI, traces of fluorescence intensity over time were created and used as substrate for subsequent analyses. Fluorescence traces were normalized to the initial fluorescence intensity (ΔF/F0). Active cells were defined as cells showing calcium waves with thresholds of 4×SDs of the baseline noise. The percentage of active cells and the amplitude of the signals for each recording were calculated and compared between each line and between the two groups (CTRL vs. SCZ).

### Matrix metalloproteinase (MMP) activity assay

To induce MMP secretion, 2×10^5^ cells/well/1mL AM medium were seeded on 12-well plates and were treated with 10 ng/ml IL-1β (Peprotech) for 24 hours. Supernatants were collected at baseline (before treatment) and following 24h incubation with IL-1β. Samples from empty wells containing only AM medium were used as negative controls. The general activity of MMPs in iPSC-astrocyte supernatants was determined using the MMP Activity Assay Kit (Abcam, Cat. no.: ab112147) following the manufacturer’s protocol. Fluorescence intensity was measured at E_x_/E_m_ = 540/590 nm using a microplate reader (GloMax, Promega, Madison, WI). MMP activity is shown in relative fluorescence units (RFU).

### Glutamate uptake assay

Indirect assessment of glutamate uptake by resting iPSC-astrocyte lines was done using a colorimetric glutamate assay kit following the manufacturer’s recommendations (Abcam; Cat. no.: ab83389). Briefly, cells were incubated in AM medium at 37°C and 5% CO_2_ for 24h then supernatants were collected and assayed to measure the amount of glutamate left in the medium. Native AM cell medium (medium control) and donor/clone-specific iPSC lines were used as controls.

### Statistics

For details about the statistical methods related to RNA-seq analysis, see the above section “Differential expression analysis” and the Supplementary Methods. *In vitro* (ELISA, MMP assay, glutamate uptake assay) results represent mean values of triplicate measurements from 5 SCZ and 5 CTRL iPSC-astrocyte cultures. Data are presented as mean ± SEM. Comparisons between groups were done by independent sample t-tests or analysis of variance (ANOVA) between groups.

## ACKNOWLEDGEMENTS

We thank all participants in this study for their invaluable contribution. In addition, the authors would like to thank Dr Erlend Strand Gardsjord who was involved in patient recruitment and clinical assessments, research nurses Eivind Bakken and Line Gundersen for obtaining, organizing and processing skin biopsies, as well as research assistants Lars Hansson, Elin Inderhaug, Evgeniia Frei, and Kristine Kjeldal for their excellent technical assistance in the cultivation of fibroblasts, and with qPCRs. The authors are grateful for the professional assistance of Dr. Zsofia Foldvari (Oslo University Hospital) in creating the artwork of Figure 1A. We acknowledge the support and use of facilities at the Norwegian Core Facility for Human Pluripotent Stem Cells, under the Norwegian Center for Stem Cell Research, part of the Regional Core Facility network of the Southeastern Regional Health Authority. We thank in particular staff members Hege Brincker Fjerdingstad and Kirstin Buchholz. The sequencing service was provided by the Norwegian Sequencing Centre (www.sequencing.uio.no), a national technology platform hosted by the University of Oslo and supported by the Functional Genomics and Infrastructure programs of the Research Council of Norway and the Southeastern Regional Health Authority. The research leading to these results has received funding from the South-Eastern Norway Regional Health Authority (#2018094) and the Research Council of Norway (#223273).

## AUTHOR CONTRIBUTIONS

Conceived and designed the experiments: A.S., I.A.A., and S.D. Performed the experiments: A.S., I.A.A., M.V. Analyzed the data: A.S., I.A.A., M.V., J.R.O., T.H., O.A.A., S.D. Contributed reagents/materials/analysis tools: J.C.G, O.A.A., S.D. All authors contributed to the writing of the manuscript.

