## Supplementary Material for "A human iPSC-astroglia neurodevelopmental model reveals divergent transcriptomic patterns in schizophrenia"

### SUPPLEMENTARY FIGURES

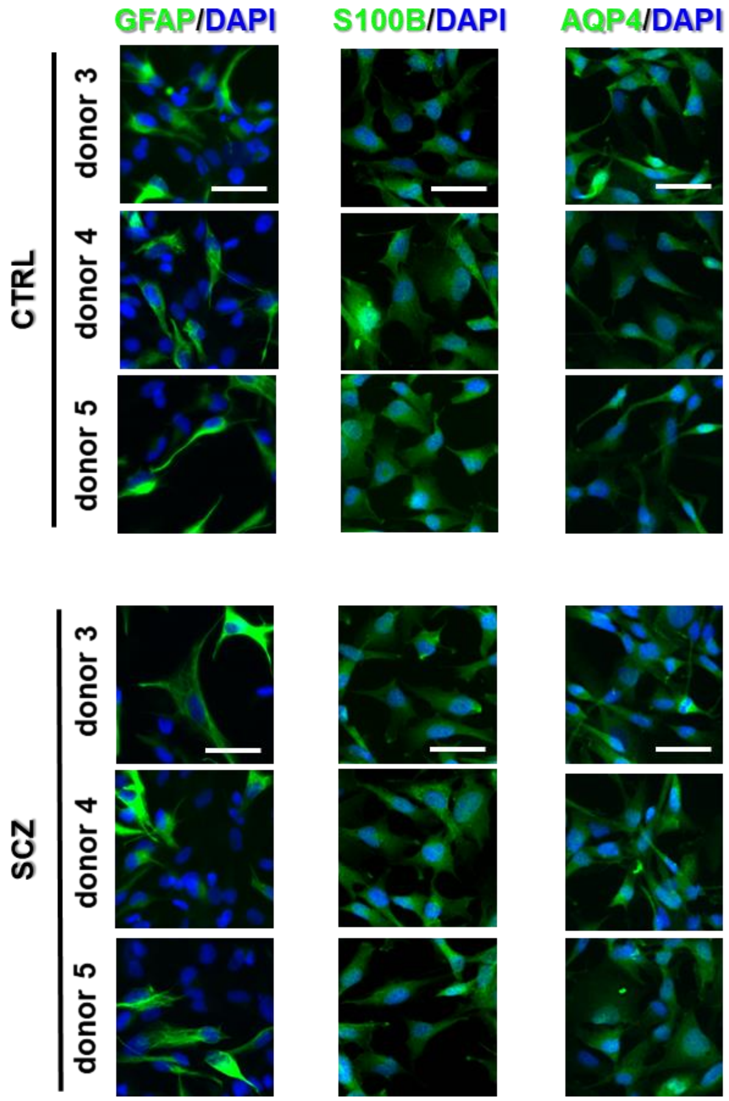


**Supplementary Figure 1. Characterization of iPSC-derived astrocytes from additional CTRL and SCZ donors (donors number 3, 4, and 5). Extended figure to Figure 1C.** Cells from day 40 cultures stained positive for the key astrocyte markers GFAP, S100B, and AQP4. Images of 3 donors with SCZ and 3 CTRLs in addition to donors 1 and 2 shown in Figure 1C. Scale bar: 200 $\mu$m.

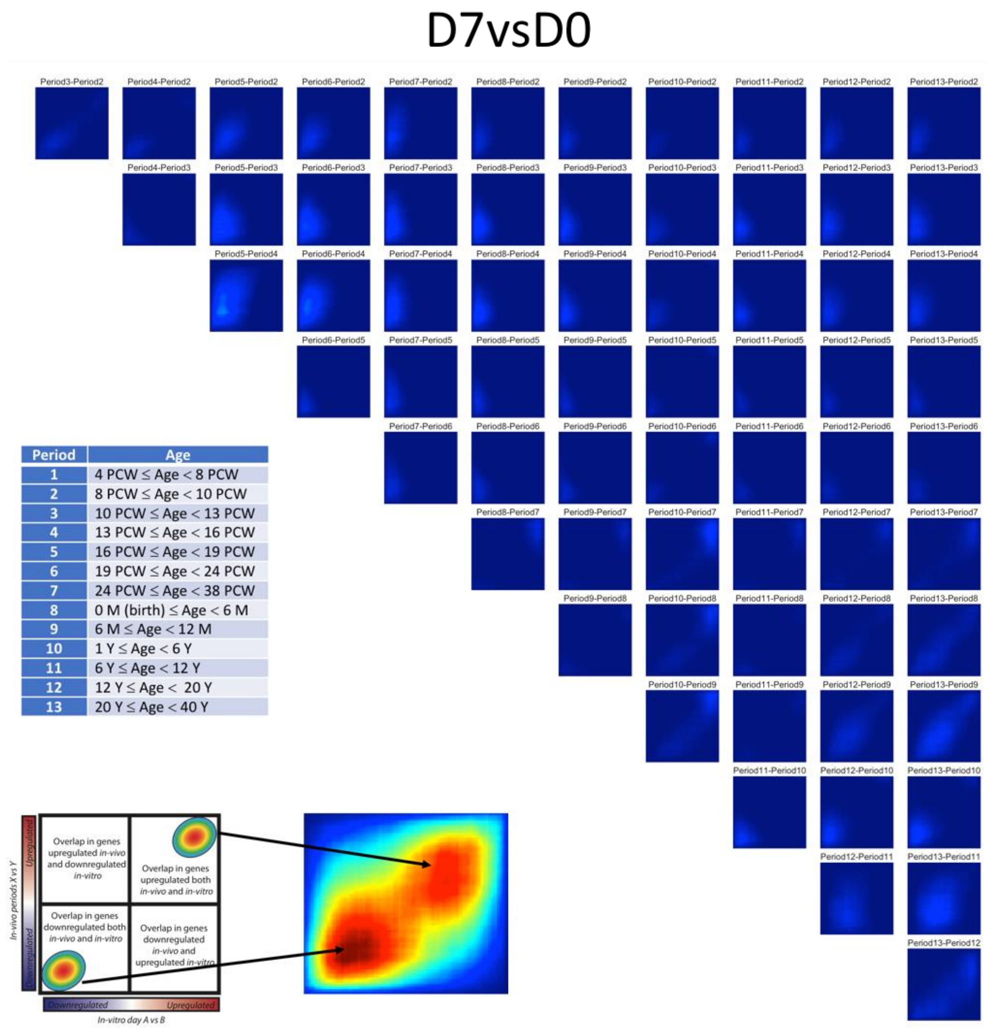


| Period | Age |
|---|---|
| 1 | 4 PCW ≤ Age < 8 PCW |
| 2 | 8 PCW ≤ Age < 10 PCW |
| 3 | 10 PCW ≤ Age < 13 PCW |
| 4 | 13 PCW ≤ Age < 16 PCW |
| 5 | 16 PCW ≤ Age < 19 PCW |
| 6 | 19 PCW ≤ Age < 24 PCW |
| 7 | 24 PCW ≤ Age < 38 PCW |
| 8 | 0 M (birth) ≤ Age < 6 M |
| 9 | 6 M ≤ Age < 12 M |
| 10 | 1 Y ≤ Age < 6 Y |
| 11 | 6 Y ≤ Age < 12 Y |
| 12 | 12 Y ≤ Age < 20 Y |
| 13 | 20 Y ≤ Age < 40 Y |

**Supplementary Figure 2.** Transition mapping between *in vitro* transition from day 0 (iPSC) to day 7 (neural rosettes) and multiple *in vivo* transitions between successive periods of brain development. The strength of correlation between *in vitro* and *in vivo* transitions is colored by -log10(p-value), see Figure 2 in the main article for color codes. Definitions of *in vivo* periods are adopted from Stein et al. [1]. The upper-right area in each map indicates overlap in genes that are up-regulated both *in vitro* and *in vivo*, while the area down to the left indicates overlap in genes that are down-regulated both *in vitro* and *in vivo*.

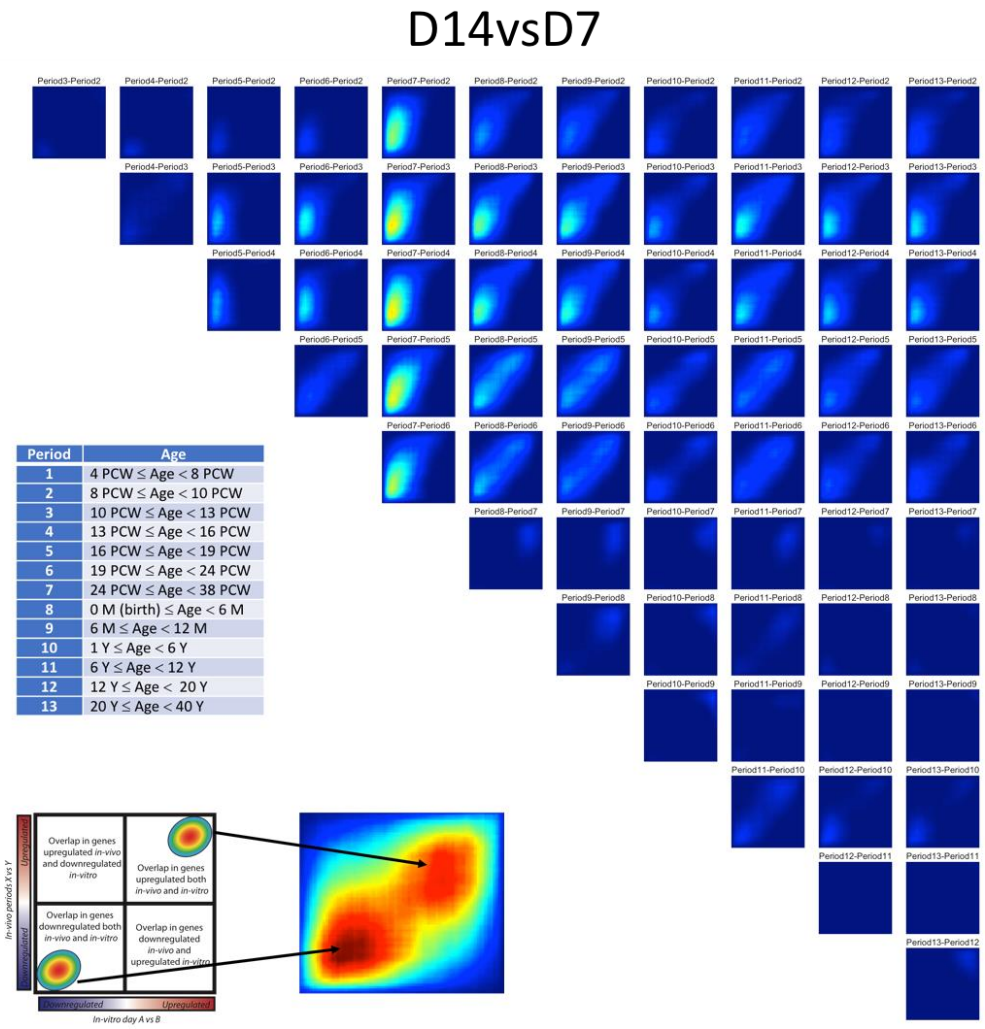


**Supplementary Figure 3.** Transition mapping between *in vitro* transition from day 7 (neural rosettes) to day 14 (neural progenitor cells) and multiple *in vivo* transitions between successive periods of brain development. The strength of correlation between *in vitro* and *in vivo* transitions is colored by -log10(p-value), see Figure 2 in the main article for color codes. Definitions of *in vivo* periods are adopted from Stein et al. [1]. The upper-right area in each map indicates overlap in genes that are up-regulated both *in vitro* and *in vivo*, while the area down to the left indicates overlap in genes that are down-regulated both *in vitro* and *in vivo*.

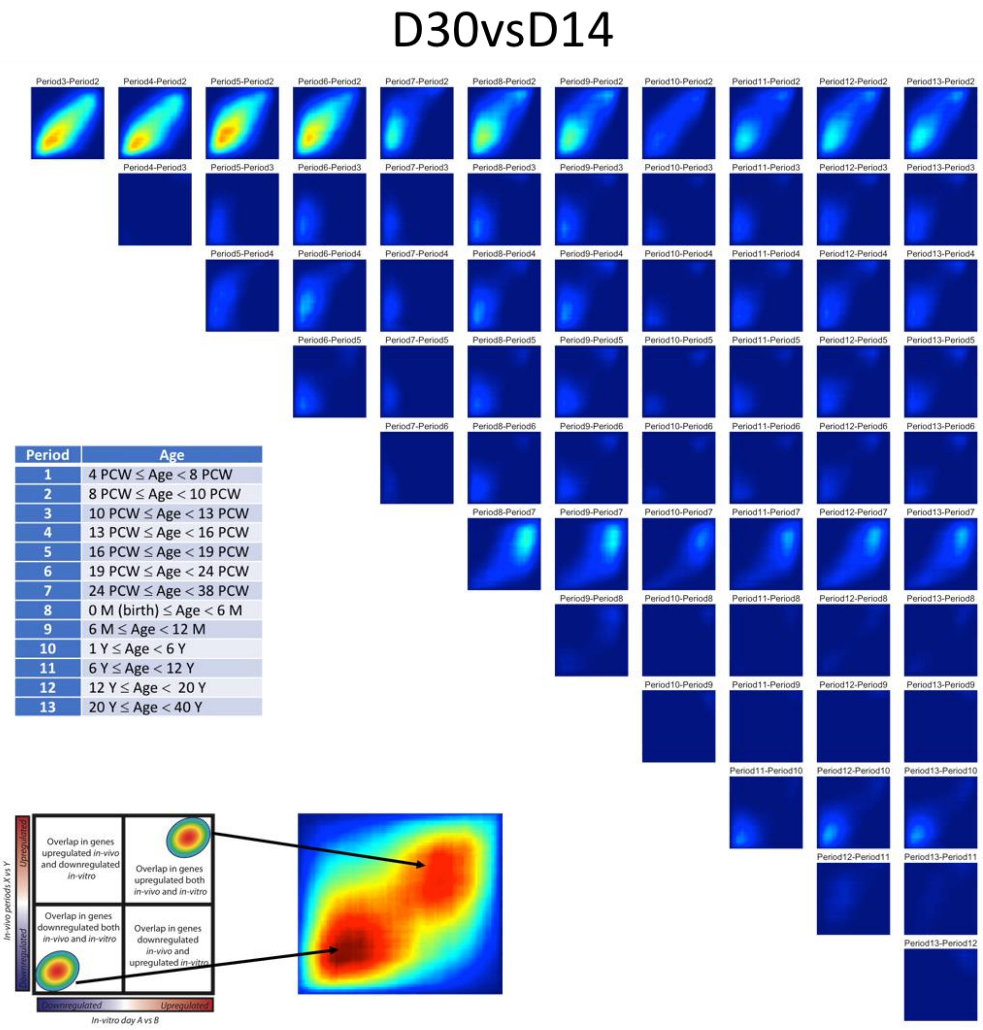


| Period | Age |
|---|---|
| 1 | 4 PCW ≤ Age < 8 PCW |
| 2 | 8 PCW ≤ Age < 10 PCW |
| 3 | 10 PCW ≤ Age < 13 PCW |
| 4 | 13 PCW ≤ Age < 16 PCW |
| 5 | 16 PCW ≤ Age < 19 PCW |
| 6 | 19 PCW ≤ Age < 24 PCW |
| 7 | 24 PCW ≤ Age < 38 PCW |
| 8 | 0 M (birth) ≤ Age < 6 M |
| 9 | 6 M ≤ Age < 12 M |
| 10 | 1 Y ≤ Age < 6 Y |
| 11 | 6 Y ≤ Age < 12 Y |
| 12 | 12 Y ≤ Age < 20 Y |
| 13 | 20 Y ≤ Age < 40 Y |

**Supplementary Figure 4.** Transition mapping between *in vitro* transition from day 14 (neural progenitor cells) to day 30 (astrocyte progenitors) and multiple *in vivo* transitions between successive periods of brain development. The strength of correlation between *in vitro* and *in vivo* transitions is colored by -log10(p-value), see Figure 2 in the main article for color codes. Definitions of *in vivo* periods are adopted from Stein et al. [1]. The upper-right area in each map indicates overlap in genes that are up-regulated both *in vitro* and *in vivo*, while the area down to the left indicates overlap in genes that are down-regulated both *in vitro* and *in vivo*.

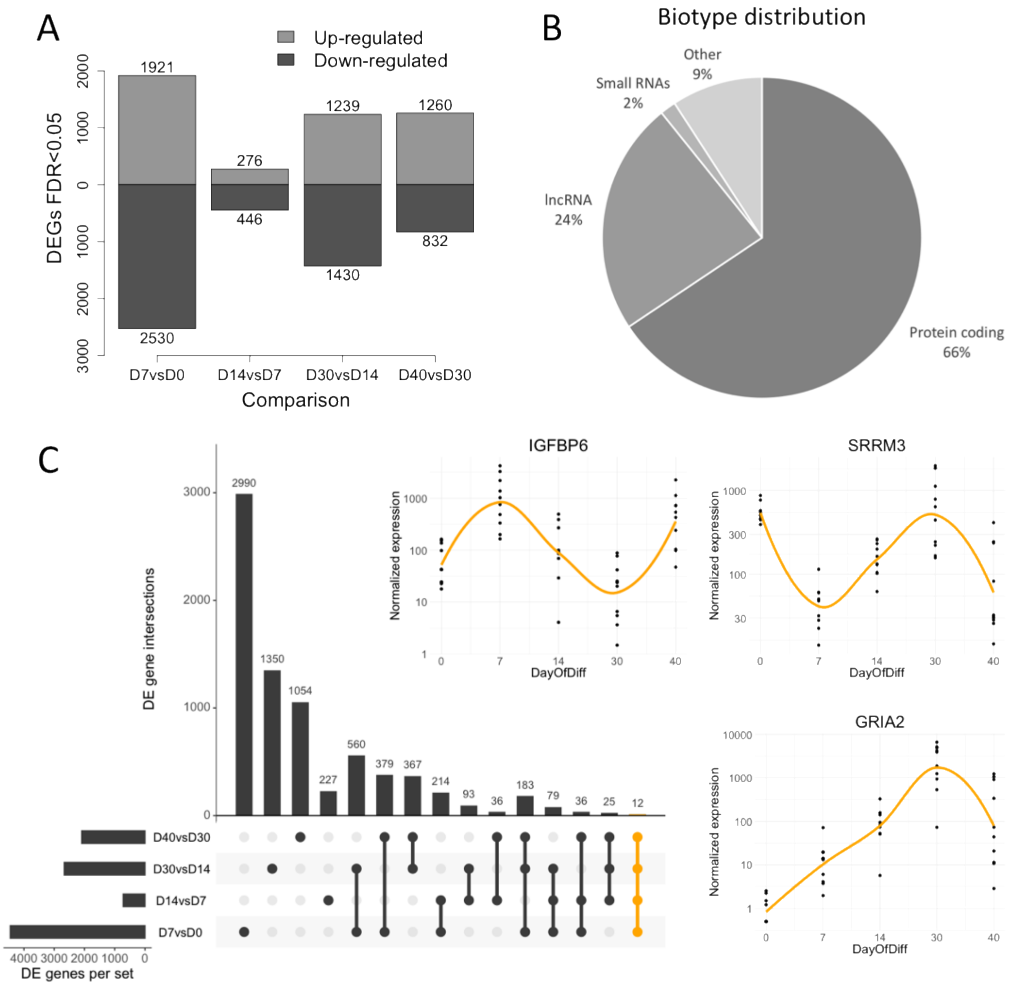


**Supplementary Figure 5. DE analyses between successive stages of *in vitro* astrocyte differentiation. A)** Barplot of identified differentiation-associated DE genes. **B)** Biotype distribution of differentiation-associated DE genes. Small RNAs include miRNAs, snRNAs, and snoRNAs. “Other” includes all remaining biotypes, including pseudogenes. LncRNA: long non-coding RNA. **C)** UpSet plot displaying gene set intersections of differentiation-associated DE genes. 12 genes were DE in all 4 comparisons (colored in orange to the right). The expression trajectories of the 3 most significant of these genes are plotted.

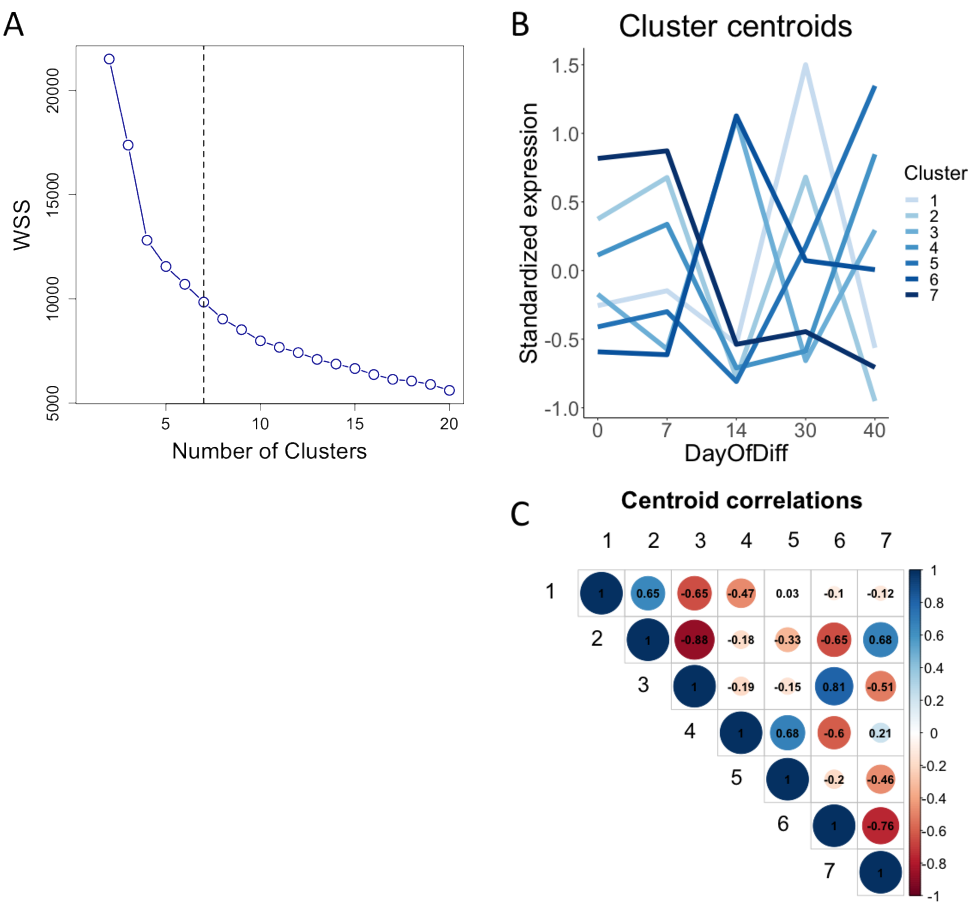


**Supplementary Figure 6. Fuzzy C-means clustering of differentiation-associated DE genes. A)** Relationship between the number of clusters and the within-cluster sum of square (WSS) value. The plot suggests that the inflection is between 5 and 10 clusters. 7 was selected as the *a priori* number of clusters to generate, as indicated by the vertical dotted line. **B)** Standardized expression profiles of cluster centroids for the 7 clusters. Centroids are colored using the same color codes as in Figure 3 in the main article. **C)** Correlogram of cluster centroids. All clusters were well separated and non-redundant, defined as a correlation of <0.85.

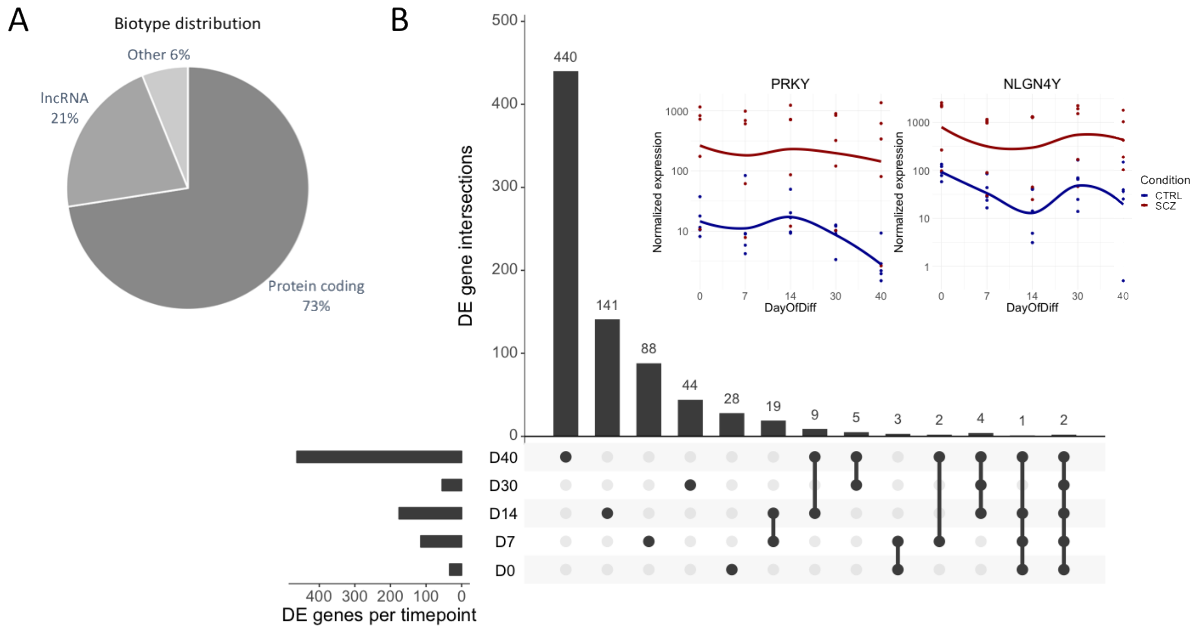


**Supplementary Figure 7. DE analyses between CTRL and SCZ samples at each stage of *in vitro* astrocyte differentiation. A)** Biotype distribution of SCZ-associated DE genes. "Other" includes miRNAs, Mt-tRNAs, and pseudogenes. LncRNA: long non-coding RNA. **C)** UpSet plot displaying gene set intersections of SCZ-associated DE genes. 2 genes were DE in all 5 comparisons. The expression trajectories of these genes are plotted.

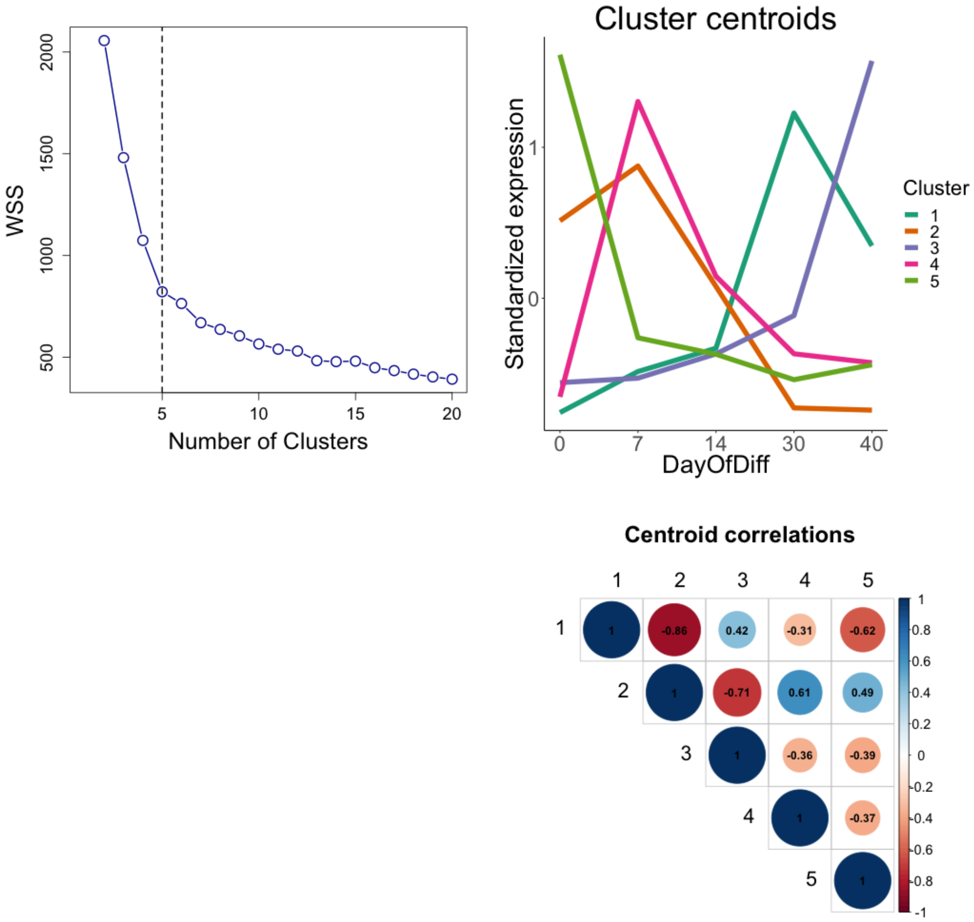


**Supplementary Figure 8. Fuzzy C-means clustering of SCZ-associated DE genes. A)** Relationship between the number of clusters and the within-cluster sum of square (WSS) value. The plot suggests that the inflection begins at 5, which was selected as the *a priori* number of clusters to generate, as indicated by the vertical dotted line. **B)** Standardized expression profiles of cluster centroids for the 5 clusters. Centroids are colored using the same color codes as in Figure 4E in the main article. **C)** Correlogram of cluster centroids. All clusters were well separated and non-redundant, defined as a correlation of <0.85.

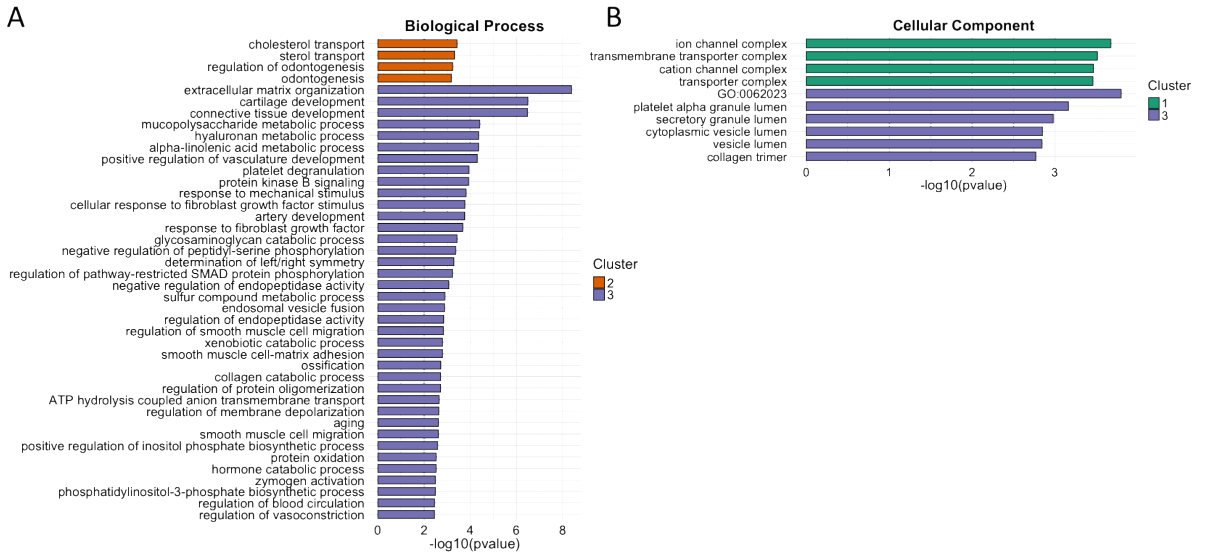


**Supplementary Figure 9. Annotations of SCZ-associated gene clusters. A)** Clusters annotated using the GO ontology “Biological Process”. Only clusters 2 and 3 were significantly enriched for one or more biological processes. **B)** Clusters annotated using the GO ontology “Cellular Component”. Only clusters 1 and 3 were significantly enriched for one or more cellular components. The length of each annotation bar indicates corresponding -log10(p-value). GO: Gene Ontology.

### SUPPLEMENTARY METHODS

#### RNA extraction and sequencing

Total RNA was extracted from the 50 samples (5 CTRL and 5 SCZ subjects at five different stages of astrocyte differentiation) using either the RNeasy Plus Mini Kit (Qiagen, Hilden, Germany) or the MagMax mirVana Total RNA Isolation Kit (Thermo Fisher) according to manufacturer's instructions. In both cases, approximately 1 million cells were used as input. RNA yield was quantified with a NanoDrop 8000 Spectrophotometer (NanoDrop Technologies, Wilmington, DE, USA) and RNA integrity was assessed with Bioanalyzer 2100 RNA 6000 Nano Kit (Agilent Technologies, Santa Clara, CA, USA). All samples had an RNA Integrity Number (RIN) above 8.5 with three exceptions: two samples at 7.8 and one sample at 6.6. Library preparation and paired-end RNA-sequencing were carried out at the Norwegian High-Throughput Sequencing Centre (www.sequencing.uio.no). Briefly, libraries were prepared with the TruSeq Stranded mRNA kit from Illumina (San Diego, CA, USA) which involves Poly-A purification to capture coding as well as several non-coding RNAs. The prepared samples were then sequenced on an Illumina HiSeq 4000 platform (Illumina) at an average depth of 50 million reads per sample using a read length of 150 base pairs and an insert size of 350 base pairs.

#### Data processing

Raw sequencing reads were quality assessed with FastQC (Babraham Institute, Cambridge, UK). To pass the initial QC check, the average Phred score of each base position across all reads had to be at least 30. Reads were further processed by cutting individual low-quality bases and removing adapter and other Illumina-specific sequences with Trimmomatic V0.32 using default parameters [2]. Since the trimming process may result in some reads being discarded and their mates thereby unpaired, only reads that remained paired after trimming were used for downstream analyses. HISAT2 [3] was then used to first build a transcriptome index based on ENSEMBL annotations, and then to map the trimmed reads to the human GRCh38 reference genome. To quantify gene expression levels, mapped reads were summarized at the gene level using featureCounts [4] guided by ENSEMBL annotations. To assess the extent to which our iPSC-derived CTRL NPCs and astrocytes resemble the cells generated by TCW et al. [5], we downloaded the raw RNA sequencing data sets from the Gene Expression Omnibus site (https://www.ncbi.nlm.nih.gov/geo/; accession number: GSE97904) and processed the data as described above.

**Differential expression analysis**

Before conducting the differential expression (DE) analyses, genes with very low to zero expression were removed by filtering out any gene with 3 or less read counts in more than 50% of the samples. DE analyses were performed using the statistical R package *DESeq2*, which provides methods to test for differentially expressed genes by use of negative binomial generalized models [6]. Two sets of DE analyses were conducted. The DE analyses in the first set were performed on two and two successive timepoints (day 7 vs. day 0; day 14 vs. day 7; day 30 vs. day 14; day 40 vs. day 30) in order to identify genes whose expression profiles change across astrocyte differentiation (differentiation-associated genes). The DE analyses in the second set were run between CTRL subjects and SCZ patients in order to identify the genes that are differentially expressed in SCZ patients across differentiation (SCZ-associated genes). Since a paired design is involved in the DE analyses in the first set (i.e. expression profiles from the same subjects are compared at two different timepoints), donor effects were included as a covariate in order to absorb subject-to-subject differences in expression at baseline within each comparison. The *DESeq2* workflow begins by taking raw read count data as input and applies an internal normalization method that corrects for sequencing depth and RNA composition. After the standard DE analysis which consists of size factor estimation, dispersion estimation, and model fitting, *DESeq2* performs an independent filtering step that optimizes the number of genes with adjusted p-values below a user-specified significance level [6]. After the pre-filtering and independent filtering steps, a total of 28,564 genes were retained for examination. In the DE analyses in the first set, a gene was considered DE if the FDR was <0.05 and the fold change >1.5. In the analyses in the second set, a gene was considered DE if the FDR was <0.10. The stricter significance criteria used in set 1 were selected in order to reduce the number of DE genes, which is typically very high when two different cell types are compared (i.e. different stages of differentiation).

**Transition mapping to cortical fetal development**

To investigate how well the *in vitro* gene expression profiles across astrocyte differentiation correspond to *in vivo* expression patterns of the developing fetal brain, we applied transition mapping (TMAP) [1] to the differentiation-associated DE genes identified in the first set of DE analyses. The TMAP method consists of serialized differential expression analyses between any two *in vitro* differentiation timepoints and any two *in vivo* developmental periods, followed by quantification of overlap using the rank-rank hypergeometric overlap test [7] for each combination of time periods. The technique generates a map of the overlap between any two systems across development. The *in vivo* developmental periods were defined from the Human Brain Transcriptome (HBT) [8], which is a spatiotemporal transcriptomic atlas generated from over 1,340 tissue samples sampled from both hemispheres of postmortem human brain specimens ranging from embryonic development to adulthood and encompassing a total of 16 brain regions. TMAP was originally developed for time course data from microarrays, but the method was subsequently modified for RNA-seq and implemented in a publicly available R script (https://github.com/dhglab/RRHO_RNAseq). The main difference between the published method and the RNA-seq-based method is the choice of the statistic used for comparison. While the published method ranks genes by p-values, the RNA-seq method uses logarithmic fold changes as this approach is more stable across different DE analysis tools. The RNA-seq method only includes cortical areas from the HBT atlas, and differential expression analyses were performed using *limma-voom* [9]. TMAP analyses across the temporal dimension of human cortical development were run for the following comparisons of *in vitro* time points: day 7 vs. day 0, day 14 vs. day 7, day 30 vs. day 14, and day 40 vs. day 30.

**Fuzzy C-means clustering**

Fuzzy C-means clustering was applied separately to the differentiation-associated and the SCZ-associated DE genes in order to cluster them according to their expression trajectory across differentiation (from day 0 (iPSC) to day 40 (astrocytes)). Fuzzy C-means clustering is a soft clustering method in which each datapoint simultaneously exists in all clusters with varying degrees of membership, where being closer to the cluster center means a higher score. Since these scores are used to position the centroids, and low scoring datapoints have a reduced impact on the position of the cluster center, fuzzy C-means clustering is more robust than hard clustering methods against the noise inherent in high-throughput profiling data such as RNA sequencing [10]. For each stage of differentiation, normalized expression values were averaged across all 10 samples (both CTRLs and SCZs) for the differentiation-associated genes and across SCZ samples for the SCZ-specific genes and then standardized. To estimate the optimal fuzzifier value, the method described by Schwämmle and Jensen [11] and wrapped into a function by Kumar and Futschik [12] was used. The number of clusters was determined by applying the within cluster sum of squared (WSS) error method as implemented in the *e1071* R package. Based on a visual expectation of the WSS plots, 7 and 5 clusters were used for the differentiation-associated and the SCZ-associated DE genes, respectively. The cluster centroids were correlated using the *corrplot* package to assess the degree of separation between the clusters and whether any of them were redundant (defined as a correlation >0.85). Only DE genes with membership >0.5 were used to generate the cluster plots and cluster annotations.

#### Cluster annotation

To annotate the 7 clusters identified for the differentiation-associated genes and the 5 clusters identified for the SCZ-associated genes. Enrichment analyses were conducted with the over-representation analysis tool *clusterProfiler* [13] using DE genes with membership >0.5 as input. Annotations were based on Gene Ontology (GO) terms [14], and a GO term was considered significantly over-represented if the FDR was <0.10.

#### Enrichment in human brain regions

To test whether the SCZ-associated DE genes were enriched in any specific brain region, we used *ABAEnrichment* [15]. This tool uses two human brain expression datasets (adult and developmental) provided by the Allen Brain Atlas [16, 17] and assigns to each anatomical structure the genes that are expressed in that structure by applying an expression threshold. It then performs ontology gene set enrichment analyses to identify significant overlaps between a custom-provided list of input genes and the genes that are expressed in a specific brain area using either a hypergeometric test or a Wilcoxon-rank sum test [15]. As cut-off for the annotation of genes to brain regions, the default 10%-steps of expression quantiles across all regions was used. The default "5_stages" dataset, which uses data from five developmental stages from the BrainSpan Atlas of the Developing Human Brain (http://brainspan.org/), was selected for the enrichment analyses. Enriched brain areas were visualized with the R package *cerebroViz* [18]. Analogous brain regions for which *ABAEnrichment* and *cerebroViz* utilize different abbreviations were named according to *cerebroViz* conventions.
